# Synthetic lipid rafts formed by cholesterol nano-patch induce early T cell activation

**DOI:** 10.1101/2023.02.22.529596

**Authors:** Yunmin Jung, Young-Joo Kim, Kunwoo Noh, Sunmi Lee, Minsuk Kwak

## Abstract

Lipid rafts cluster at the immunological synapse during activation and serve as signaling hubs in T cells. While cholesterol plays a crucial role in lipid raft formation, the impact of their spatial configuration remains less understood. Here, we introduce programmable DNA origami, cholesterol nano-patch, for nanoscale spatial control of cholesterols on live T cell membranes, enabling us to elucidate their roles in lipid raft formation, receptor activation, and intracellular signaling. We demonstrate that CNPs with high-density cholesterol arrangements efficiently bind to the T cell plasma membrane and form a large and polarized coalescence, leading to the stabilization of ordered lipid membrane domains by increasing the local concentration of cholesterols. These synthetic lipid rafts colocalize with flotillin-1, a raft-associated protein, and promote the membrane reorganization and physical segregation of key signaling molecules, such as T cell receptors, Lck and LAT kinases, and CD45. Consequently, this leads to early T cell activation in the absence of antigenic stimulation. Our DNA origami approach demonstrates the direct role of cholesterol in lipid raft formation, leading to membrane phase separation and protein reorganization, which are sufficient to induce early T cell activation.

## Introduction

Live cell membrane contains small and dynamic nanodomains enriched in cholesterol and sphingolipids known as lipid rafts ^1,2^. Due to their more ordered, thicker, and less fluidic characteristics, lipid rafts contribute to the spatial organization of membrane proteins and influence broad arrays of cellular signaling events ^1,3,4^. Nano-sized lipid rafts (10-200nm) can be clustered by protein-protein or protein-lipid interactions to form large (>300nm) domains serving as cell signaling platforms ^1,3,4^. Such lipid rafts clustering and accumulation has been observed at immunological synapse during T cell activation, serving as signaling platforms by driving spatial reorganization of key signaling molecules ^5–8^, including the recruitment of TCR and activating kinases such as Lck, LAT, and the exclusion of CD45^9–11^.

Liquid-liquid phase separation (LLPS) of lipid molecules within the membrane is a fundamental physical property that creates heterogeneous compartments on the cell membrane ^3,12,13^. Cholesterol is a key component that induce liquid ordered (Lo) – liquid disordered (Ld) phase separation of plasma membrane ^14,15^. This Lo-Ld phase separation can be attributed to the favorable interactions between cholesterol and saturated acyl chains and the lateral immiscibility of cholesterol with unsaturated acyl chains ^16–18^. The longer, more saturated acyl chains in the Lo domain allow lipids to pack more tightly and form a thicker and less fluidic membrane that influences the spatial organization of membrane proteins and cell function ^3,9,19,20^. While cholesterol-driven large-scale LLPS has been observed in model membranes at their thermodynamic equilibrium ^14,15^, such observation and investigation of LLPS induced by cholesterol in the non-equilibrium cell membrane remains challenging ^13^.

The role of cholesterol in T cell activation has been suggested that cholesterol can either promote or suppress T cell signaling by inducing TCR nanoclustering ^21^ or stabilizing the TCR inactive conformation, respectively ^22,23^. Global alteration of cholesterol levels through the treatment of squalene or methyl-β-cyclodextrin has demonstrated a profound influence on cholesterol homeostasis in T cell functions ^24–27^. However, the effect of local cholesterol concentration on the liquid-liquid phase separation (LLPS) of the plasma membrane in live cells and the lipid raft-driven protein reorganization in T cells remains poorly understood due to methodological limitations in controlling the spatial distribution of cholesterol.

DNA origami is a molecular fabrication technique enabling the construction of a customized nanostructure with desired shape and size, and it allows facile manipulation of spatial configuration of biomolecules with nanoscale precision ^28,29^. To directly investigate the roles of cholesterol in the coalescence of lipid rafts and protein partitioning in T cells, we designed a cholesterol-conjugated DNA origami nanostructure (termed as cholesterol nano-patch or CNP), as a nano-sized lipid raft-like monomer when it attaches to the membrane. CNPs enable precise patterning and display of cholesterol molecules in a programmable manner. We demonstrated that CNPs bind to the outer leaflet of the T cell membrane and form a large and stably polarized cluster. We defined this large coalescence of multiple CNPs segregated into a distinctive membrane domain via enrichment of cholesterols as a synthetic lipid raft. We showed that the high density of cholesterols on CNPs is more effective in the formation of synthetic lipid rafts as it increases the local concentration of cholesterols on live T cell membranes. Using the CNP platform, we investigated how the nanoscale arrangement of cholesterols in regulating the lipid raft formation, spatial reorganization of membrane proteins, and T cell signaling responses.

## Results

### Design of DNA origami structure for cholesterol patterning

We fabricated a cholesterol-attached sheet-like DNA nanostructure, CNP, employing a DNA origami method. This approach allowed us to create a molecular template where the number or arrangement of cholesterols can be precisely programmed (**Fig. 1**). By hybridizing multiple short single-stranded DNAs (ssDNAs) with a long ssDNA strand and folding it into a desired geometry, we achieved a precise shape for the resulting DNA nano-patch (NP), a square sheet measuring approximately 60 nm on each side and 6 nm in height. Cy5-fluorophores were placed at each of its four corners for visualization (**Fig. 1A** and **Table S1**). In order to achieve site-specific anchoring and precise patterning of cholesterols, an array of binding sites was designed on one side of NP. On these binding sites, ssDNAs (DNA handles) could form complexes with cholesterol-conjugated DNAs (Chol-DNAs) and assist DNAs (**Fig. 1A**). Each NP could display up to 72 Chol-DNAs in an array of 12 rows and 6 columns, with a gap of 3.5 nm and 7 nm, respectively. This arrangement allowed for an inter-cholesterol spacing ranging from 3.5 to 38.5 nm (**Fig. 1A,B**). Cholesterols on CNPs facilitated hydrophobic interactions with the lipid bilayers of plasma membranes (**Fig. 1C**). The estimated height of the hybridized Chol-DNA is 10.9 nm (32 base-pairs). To control the local concentration of cholesterols, we varied their spatial distribution by patterning 18- (0.005 cholesterols/nm^2^), 36- (0.01 cholesterol/nm^2^), or 72- (0.02 cholesterols/nm^2^) cholesterols on a NP, termed 18-CNP, 36-CNP, and 72-CNP, respectively (**Fig. 1B**). In order to test the efficient control of the number of desired conjugates, we substituted the fluorescent dye (Cy5) for cholesterol in CNPs and measured the fluorescent intensities of the conjugated NPs. The results showed that the intensities increased linearly depending on the available binding sites on NPs (**Fig. S1A**). We also confirmed the proper folding of the structure and the high assembly yield of the NPs through agarose gel electrophoresis (**Fig. S1B**) and transmission electron microscope (TEM) imaging (**Fig. 1D**). The purified NPs remained stable in the RPMI buffer for 30 minutes, consistent with our CNP binding conditions for cells (**Fig. S1C,D**). Furthermore, we confirmed the ability of binding of CNPs to the surface of synthetic small lipid vesicles using TEM imaging (**Fig. 1E**), demonstrating their efficient interaction with the lipid membrane.

**Figure 1.**
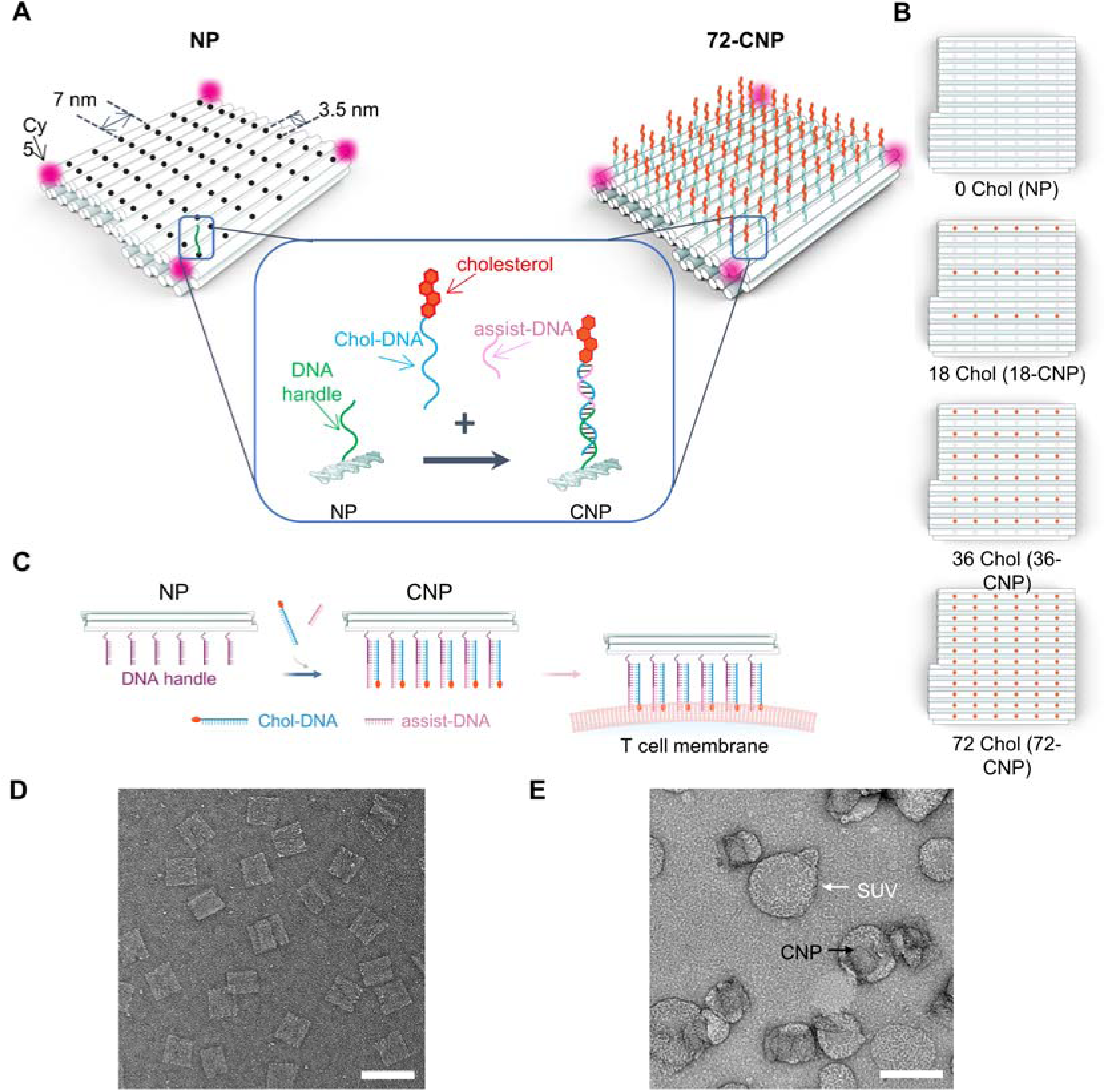
Cholesterol nano-patch (CNP) displaying multiple cholesterols. **(A)** Schematic illustrations for a DNA nano-patch (NP, upper left) and a CNP structure containing 72 cholesterols (72-CNP, upper right). DNA-NP is a square DNA origami pegboard with dimensions of 60 x 60 nm with a 6 nm height and each cylinder represents a double-stranded DNA (dsDNA). Cy5-conjugated single-stranded DNAs (ssDNA) are attached at the four corners to introduce the fluorescence imaging capacity. The black circles on the NP separated by 3.5 and 7 nm in rows and columns, respectively, represent potential locations where the DNA handles can be incorporated for attachment of Chol-DNAs. Chol-DNAs are complexed to the DNA handles presented on the DNA NP *via* Watson-Crick oligonucleotide hybridization. Assist-DNA strands are added together to form a stable duplex form. **(B)** Schematic illustrations of the spatial arrangement of cholesterol molecules in 18-CNP, 36-CNP, or 72-CNP. The positions of cholesterols are marked with orange dots. **(C)** Schematic illustrations showing the conjugation strategy of CNPs onto T cell membrane. First, CNPs are prepared by complexing Chol-DNAs and assist DNAs with DNA handles presented on the NPs. Then, CNPs bind to the membranes via hydrophobic interactions between cholesterol and lipid bilayer. **(D**-**E)** Representative TEM images of DNA-NPs (D) and CNPs bound on the surface of small unilamellar vesicles (SUVs) (E). Scale bars in (D) and (E) are 100 nm.

### Formation of synthetic lipid rafts on T cell membranes

We examined whether the CNPs can drive the formation of lipid rafts by increasing the local concentration of cholesterol on T cell membranes. We employed two different binding strategies for CNP attachment to plasma membranes: First, Chol-DNA, assist-DNA, and DNA NP was sequentially applied to Jurkat T cells (**Fig. S2A,B**); Second, CNPs were pre-assembled in a tube and added to T cells as a single-step reaction (**Fig. 1C**; **Fig. 2A**). The mean fluorescent intensity (MFI) of single cell-bound CNPs was 3.4x higher in the single step binding reaction than in the sequential reaction, indicating the direct addition of the pre-assembled CNPs to cells was more efficient than the sequential binding reaction (**Fig. S2C).** The CNPs binding induced spontaneous formation of micron-sized (∼14.3 ± 5 µm in perimeter) polarized clusters, synthetic lipid rafts, on the T cell membranes (**Fig. 2A; Fig. S2B**). Next, to investigate the impact of the nanoscale distribution of cholesterol, we applied CNPs with varying numbers of attached cholesterols (**Fig. 1B**). The control NPs without cholesterol (0-chol) did not bind to the Jurkat T cell membranes. CNPs with a higher number of cholesterols exhibited increasingly higher efficiency in forming polarized CNP clusters on the membrane (**Fig. 2B,C**). In 72-CNP-treated samples, 69% of cells formed single-polarized synthetic lipid rafts on their membranes, and the percentages of this single-polarized population dropped to 50% and 33% in 36-CNP- and 18-CNP-treated cells, respectively (**Fig. 2C; Fig. S2D**). We also found that a minority of cells (< 5%) had more than one polarized cluster (**Fig. 2C; Fig. S2E**). Next, we assessed the stability of CNP-induced synthetic lipid rafts under standard cell culture conditions and their effects on T cell proliferation. We observed that T cells treated with 72-CNPs maintained the polarized CNP caps for 4 hours, while a majority of the 18- and 36-CNPs became detached from the cells (**Fig. 2D; Fig. S3**). The binding of CNPs did not affect cell proliferation for 4 days, regardless of the number of cholesterols on the CNPs (**Fig. S4)**. The 72-CNPs similarly induced the formation of the synthetic lipid rafts in freshly isolated primary mouse and human T cells (**Fig. 2E**) as well as in differentiated effector T cells **(Fig. S5A**), and these synthetic lipid rafts remained almost intact for 4 hours (**Fig. S5B,C**). Together, these results suggest that cholesterol-mediated binding of CNPs drives membrane segregation, and a tighter cholesterol distribution on CNPs enhances membrane binding efficiency, synthetic lipid raft formation, and stability.

**Figure 2.**
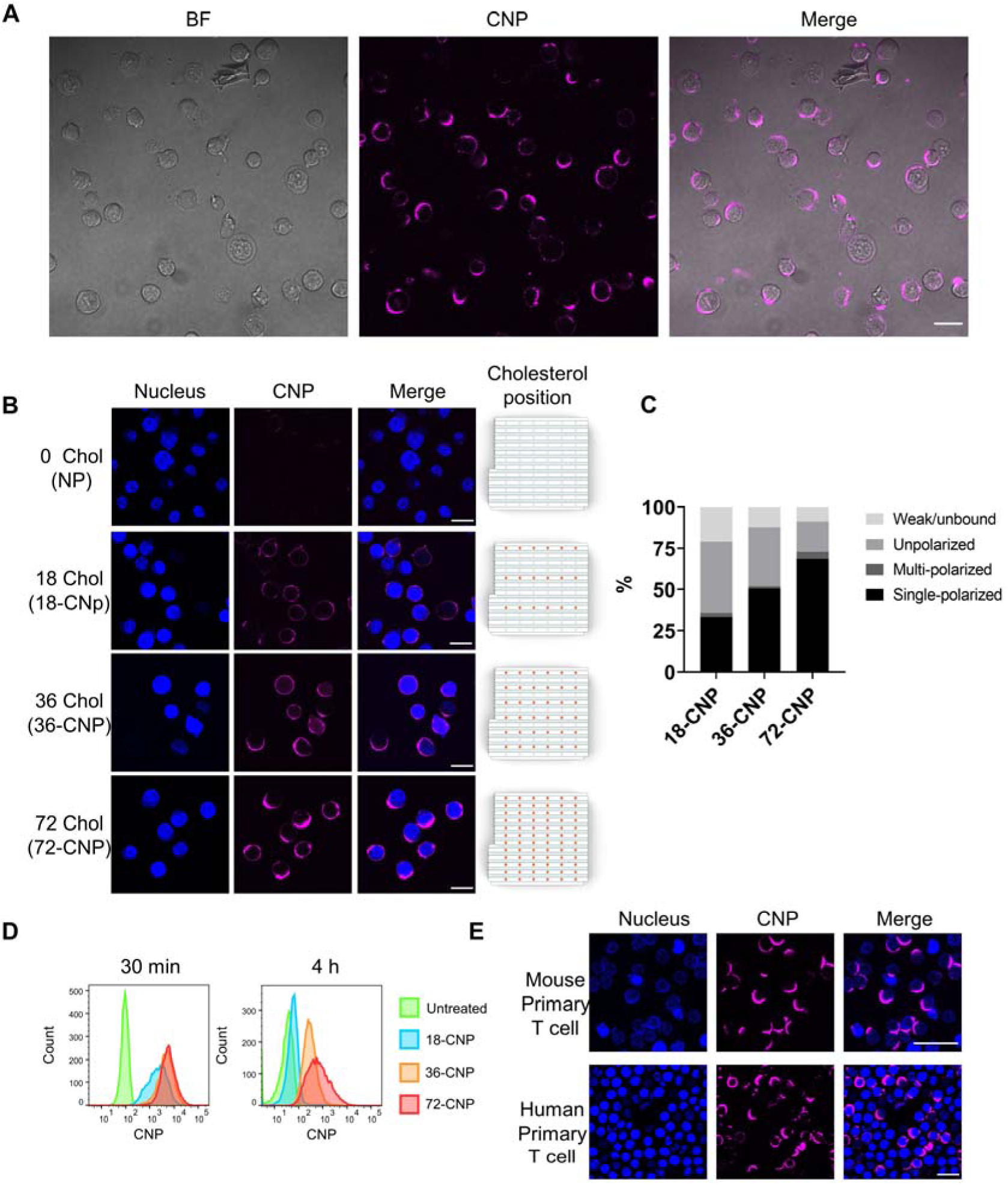
Synthetic lipid rafts on T cell membranes. **(A)** Representative confocal images of Jurkat T cells incubated with 72-CNP for 30 min. Bright-field (BF) (gray, left), 72-CNP (magenta, middle), and the merged images (right) are shown. **(B)** Effects of increasing the number of cholesterols on the formation of the synthetic lipid rafts. Representative confocal images of Jurkat T cells incubated with DNA NPs without cholesterol or complexed with 18- (18-CNP), 36- (36-CNP), or 72- (72-CNP) cholesterols for 30 min. Images of nuclei stained with Hoechst (blue, left), CNP (magenta, middle left), and merged images (middle right) for each sample are presented. The orange dots on the schematic drawings (right) indicate the positions of cholesterols. **(C)** The percentage of weak/unbound (light gray), unpolarized (gray), multi-polarized (dark gray), or single-polarized (black) synthetic lipid raft positive cells were counted from Jurkat T cells incubated with 18-(n=120), 36-(n=78), or 72-CNP (n=89) for 30 min (see the **Fig. S2D-G**). **(D)** Intensity distributions of the different CNPs on single cells incubated for 30 min (left) or 4 h (right) were analyzed by flow cytometry. **(E)** Representative confocal images of freshly isolated mouse T cells (upper) or human T cells incubated with 72-CNP for 30 min. Scale bars in (A), (B) and (E) are 20 µm. All data were collected from at least three independent experiments.

### Synthetic lipid rafts stabilize more ordered phase of the plasma membrane

We then investigated whether the synthetic lipid rafts induced by CNPs influence the lipid phases (Lo-Ld) within the plasma membrane of T cells. To quantify the membrane lipid fluidity and order, we labeled the 72-CNP bound T cells with C-laurdan, a polarity-sensitive fluorescent probe (**Fig. 3A**) ^30^. The emission spectra of C-laurdan shift towards shorter wavelengths in the Lo phase environment compared to the Ld phase lipid environment. As a result, the ratiometric calculation of fluorescence intensity between the 400-460nm and 470-530nm emission channels, known as generalized polarization (GP), becomes higher in the Lo phase than in the Ld phase ^31^. We compared the GP values between CNP-bound (raft) and unbound (non-raft) regions of the plasma membrane, determined using an automated segmentation imaging process (**Fig. 3B**). Remarkably, the GP values in the raft exhibited a significant increase compared to the non-raft region of the plasma membrane (**Fig. 3B-D**). The mean GP difference between the CNP-bound and non-bound regions for 51 cells was 0.081 (**Fig. 3D**), which closely resembles the GP change of 0.1-0.15 observed in the presence of cholesterol in the synthetic phospholipid membrane ^32^. This result indicates that CNP segregation on the membrane stabilizes Lo-domain of lipid domain, consequently causing microscale phase separation of the plasma membrane of T cells.

**Figure 3.**
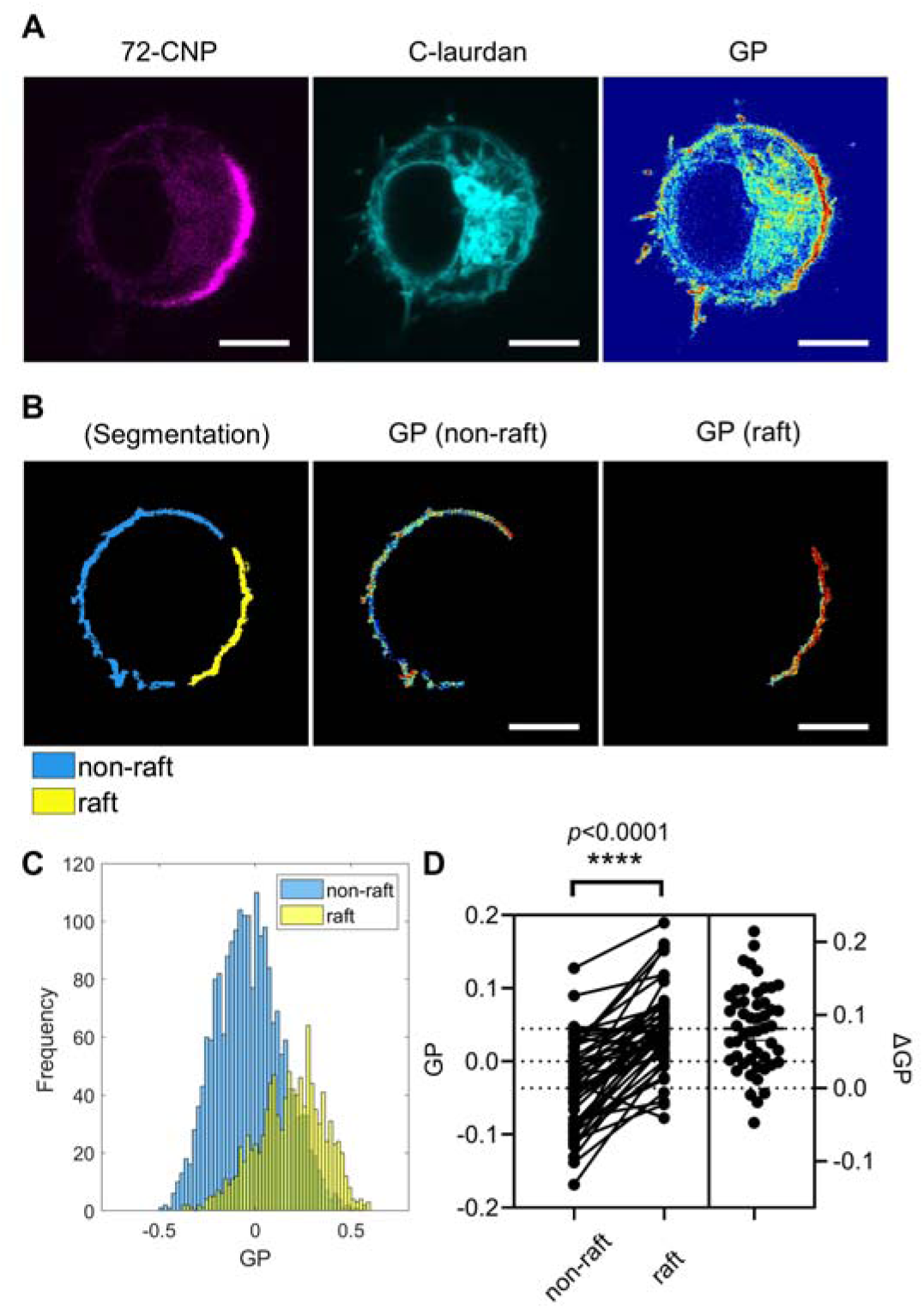
Phase separation of plasma membrane of a live T cell induced by CNPs. **(A)** Representative confocal images of a Jurkat T cell with CNPs (magenta), C-laurdan (cyan), and GP imaging **(B)** GP images of the segmented plasma membrane (left), the CNP-unbound area (non-raft, middle), and the CNP-bound region (raft, right) from the cell in (A). **(C)** The distribution of GP values within the non-raft and raft regions in (B). **(D)** Median GP values of the non-raft and raft regions were measured from 51 cells collected from three independent experiments (left panel). Each set of two connected dots represents data from a single cell. The right panel displays the distribution of differences in GP (ΔGP). Each dot representing data from a single cell. p-values (****, p ≤ 0.0001) were calculated using a two-tailed paired t-test. Color bars indicate the GP values. Scale bars in (A) and (B): 5µm.

### Synthetic lipid rafts enhance and stabilize the polarization of flotillin-1

We further explored the nature of the CNP-induced synthetic lipid rafts on T cell membrane by monitoring the spatial organization and dynamics of the CNPs and a lipid raft associated protein, flotillin-1 (Flot1) ^33,34^. Flot1 forms multimeric complexes with flotillin-2 (Flot2), and these complexes are anchored to the inner leaflet of the membrane through palmitoylation or myristoylation, leading to their segregation into lipid rafts ^35,36^. In T cells, Flot1 is weakly pre-polarized on the membrane in the resting state, and this polarization is significantly enhanced upon stimulations ^37–41^. Here, we created synthetic lipid rafts using the 72-CNPs on Jurkat T cells that stably express Flot1 with Halo-tag (Flot1-Halo). Using live cell time-lapse imaging, we monitored the dynamics of Flot1 for 30 minutes after adding the CNPs. As reported previously, we observed cap-like preassembled Flot1 in the membrane (**Fig. 4A-D**), which were more strongly polarized upon 72-CNPs addition (50%) than untreated controls (32.8%) (**Fig. S6A**). CNP-induced synthetic lipid rafts exhibited significant colocalization with Flot1-caps (95% of the Flot1-polarized cells (n=133)) after 30 minutes incubation (**Fig. 4A**). To quantify the level of augmentation of Flot1 at the CNP-bound synthetic lipid raft, we segmented sub-areas of the cell membrane into CNP-bound (raft, yellow) and unbound (non-raft, light blue) regions of the plasma membrane (**Fig. 4B**) and calculated the normalized Flot1 intensities between these segmented regions in each cell. The average of the median Flot1 intensities within the raft regions increases 3 times more than that in the non-raft regions across 59 cells (**Fig.4C**). The substantial colocalization efficiencies suggest that CNP-mediated synthetic lipid rafts can effectively recruit Flot1, thereby mimicking the coalescence of native lipid rafts. (**Fig. 4D**). Using live cell time-lapse imaging, we identified three distinct outcomes: (1) Initially, CNPs bound homogeneously to Flot1-unpolarized cells and induced co-clustering of Flot1-caps and CNP clusters (40 %) (**Fig. S6B**; **Movie S1**); (2) CNPs initially bound homogeneously to Flot1-polarized cells and subsequently moved toward the Flot1 caps (33 %) (**Fig. S6C; Movie S2**); and (3) CNPs preferentially bound to the weakly polarized Flot1 caps and enhanced their polarization (27 %) (**Fig. S6D; Movie S3**). Overall, CNP-mediated synthetic lipid rafts exhibited strong colocalization with Flot1 caps in T cells, and the binding of CNPs on the membrane further enhanced the polarization of Flot1, resembling previous observations where T cells displayed augmented lipid raft formation upon activation by anti-CD3/CD28, chemokines, or lipid raft crosslinking ^39–41^.

**Figure 4.**
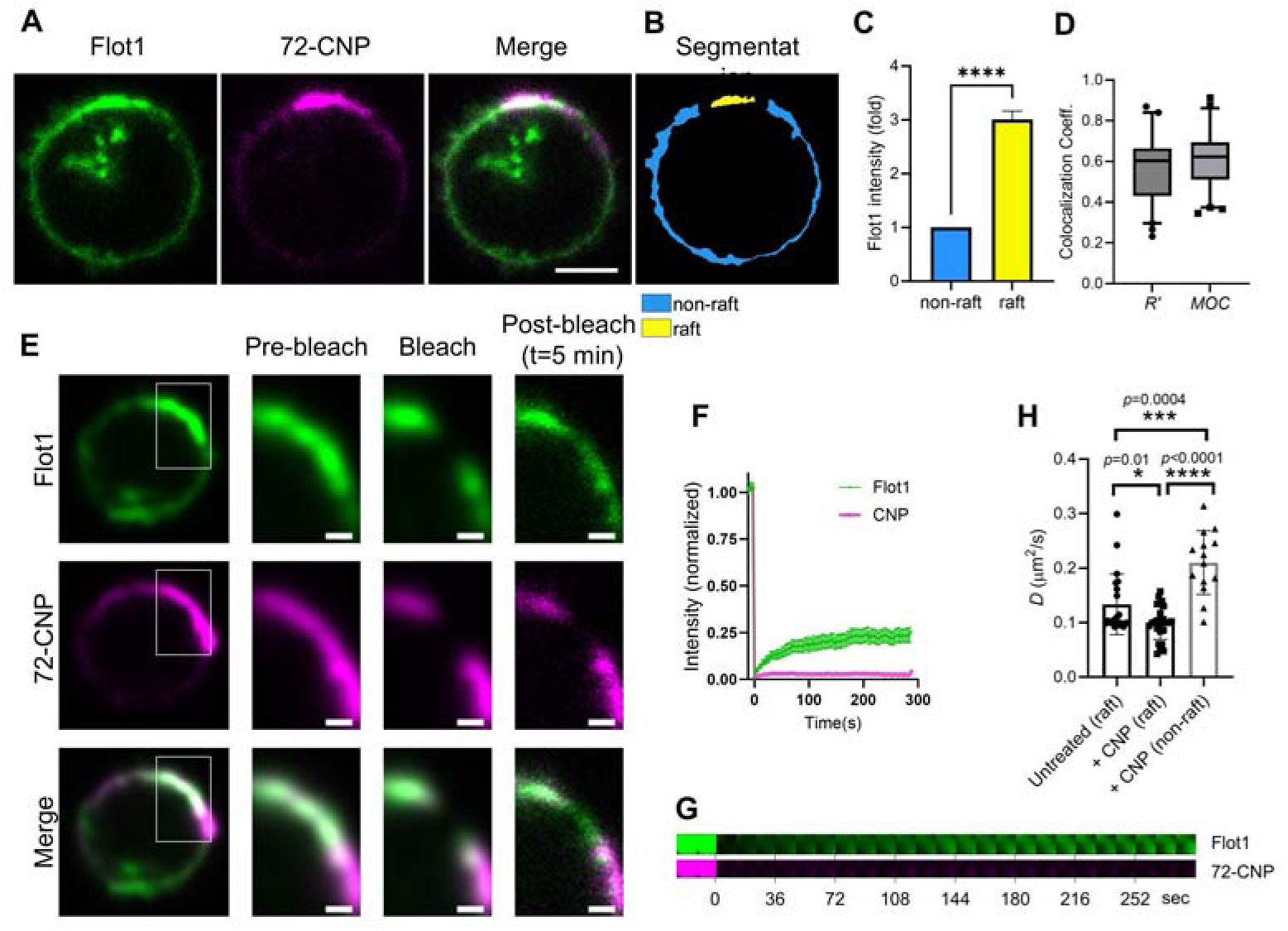
Co-polarization of CNPs-induced synthetic lipid raft and the raft-associated protein Flot1. **(A)** Representative confocal images of a Jurkat T cell stably expressing Flot1-Halo incubated with 72-CNP for 30 min. Flot1-Halo labeled with Janelia 549 Halo-tag ligand (green, left), 72-CNP (magenta, middle), and the merged images (right) are presented. **(B)** The segmented polarized CNP-bound (raft, yellow) and unbound (non-raft, light blue) regions of the plasma membrane of the cell displayed in (A)**. (C)** The means of the median Flot1 intensities corresponding to the segmented non-raft (light blue) and raft (yellow) regions were calculated (n=59 cells collected from three independent experiments). The median intensities for each regions were normalized to the median non-raft intensity of each cell. **(D)** Colocalization coefficients (Pearson’s correlation coefficient (*R’*) and Manders overlap coefficient (*MOC*)) between Flot1 and 72-CNP within the segmented regions of the data set in (C) were calculated. The lines within the boxes represent the means and the dots above and below the whiskers represent the outliers that are either greater than 95th or less than 5th percentile. **(E-G)** FRAP analysis of the lateral mobility of Flot1 and 72-CNP bound on the membrane. Representative time-lapse confocal images of Flot1 (green) and 72-CNP (magenta) in the area marked by a square in the left images taken before and after bleaching (E). The mean FRAP tracings of Flot1 (green) and 72-CNP (magenta) after photobleaching for 5 minutes from live Jurkat T cells treated with 72-CNP. The data represent means ± SEM (n = 18 cells from three independent experiments) (F). The filmstrips showed fluorescence recovery of the bleached area of the cell in (E) marked in **Movie S5** (G). **(H)** The mean diffusion coefficients (*D*) of Flot1 were analyzed in untreated cells (left bar, n=21) at the raft region (where Flot1 is polarized), and in 72-CNP treated cells, both at the raft (middle, n=26) and non-raft (right, n=14) areas, using FRAP. Each dot represents data collected from a single cell, and the data are presented as the mean ± SD of all cells pooled from three independent experiments. *p*-values (*, *p* ≤ 0.05; ***, *p* ≤ 0.001, ****, *p* ≤ 0.0001) were calculated using a two-tailed unpaired t-test. Scale bars in (A): 5µm; in (E): 1µm.

Next, we monitored the lateral mobility of 72-CNPs on the Jurkat T cell membrane using fluorescence recovery after photobleaching (FRAP) ^42^. Interestingly, there was almost no detectable recovery of 72-CNP fluorescence in the synthetic lipid rafts for 5 minutes after rapid photobleaching, whereas Flot1 fluorescence in the synthetic lipid raft recovered slowly (**Fig. 4E-H; Fig. S7; Movie S4; Movie S5).** The slower and partial recovery of the polarized Flot1 at the CNP-bound (raft) region is significantly different from the Flot1 located in the non-raft region of the membrane (**Fig. S7D**). This result is similar to a previous report on the dynamics of polarized versus non-polarized flotillins ^39^. The diffusion coefficient (*D*) of Flot1 in CNP-unbound (non-raft) areas was ∼2.1 times higher than *D* of Flot1 in the CNP-bound (raft) areas (**Fig. 4H**). Notably, there was a significant decrease in *D* for Flot1 in the synthetic lipid rafts induced by CNP binding compared to that in the naturally formed polarized area (raft) (**Fig. 4H**). These results demonstrate that the formation of highly stable CNP clusters on the outer leaflets affects the stabilization of more ordered lipids on the inner leaflets of the membrane. Overall, CNP-mediated synthetic lipid rafts exhibited strong colocalization with polarized Flot1-preassembles in T cells, and the binding of CNPs on the membrane further enhanced the polarization of Flot1 and stabilizes its motility, resembling previous observations where T cells displayed augmented lipid raft formation upon activation by anti-CD3/CD28, chemokines, or lipid raft crosslinking ^39–41^. These results support that CNP-induced synthetic lipid rafts exhibit the characteristics of natural lipid rafts and influence the dynamics of raft-associated proteins on the membrane.

### Synthetic lipid rafts reorganize membrane proteins and induce early T cell activation

Next, we investigated whether CNP-mediated synthetic lipid rafts can influence the rearrangement of membrane proteins involved in the early signaling of T cell activation. We first assessed the distribution of CD45 on the membrane after CNP binding. CD45 exclusion from the immunological synapse, resulting in decreased phosphatase activity in that region, is a crucial step in T cell activation ^43–46^. Previous reports have indicated that artificially induced CD45 exclusion can induce T cell activation even in the absence of antigen ^44^. A recent study revealed that CD45 is excluded from the microvilli tips due to the ticker membrane formed by cholesterol, prior to contact with antigen-presenting cells (APCs) ^47^. We investigated whether CD45 is segregated from the CNP-bound synthetic lipid rafts. Upon the addition of 72-CNPs, CD45 molecules partitioned away from the synthetic lipid rafts in 84% of CNP-polarized cells (n=104) (**Fig. 5A**). To quantify the level of exclusion, we segmented sub-areas of the cell membrane into CNP-bound (raft, marked in yellow) and unbound (non-raft, marked in light blue) regions of the plasma membrane (**Fig. 5B**) and calculated the normalized CD45 intensities between these segmented regions in each cell. The average of the median intensities of CD45 within the raft regions was only 33% of the CD45 intensities in non-raft regions across 51 cells (**Fig. 5C**). The negative Pearson’s correlation coefficient (*R’ = - 0.57±0.09*) between the CD45 and 72-CNPs within the segmented membrane regions, coupled with the near-zero Manders overlap coefficient (*MOC = 0.04±0.03*), indicates mutual segregation between the CD45 and 72-CNPs (**Fig. 5D**).

**Figure 5.**
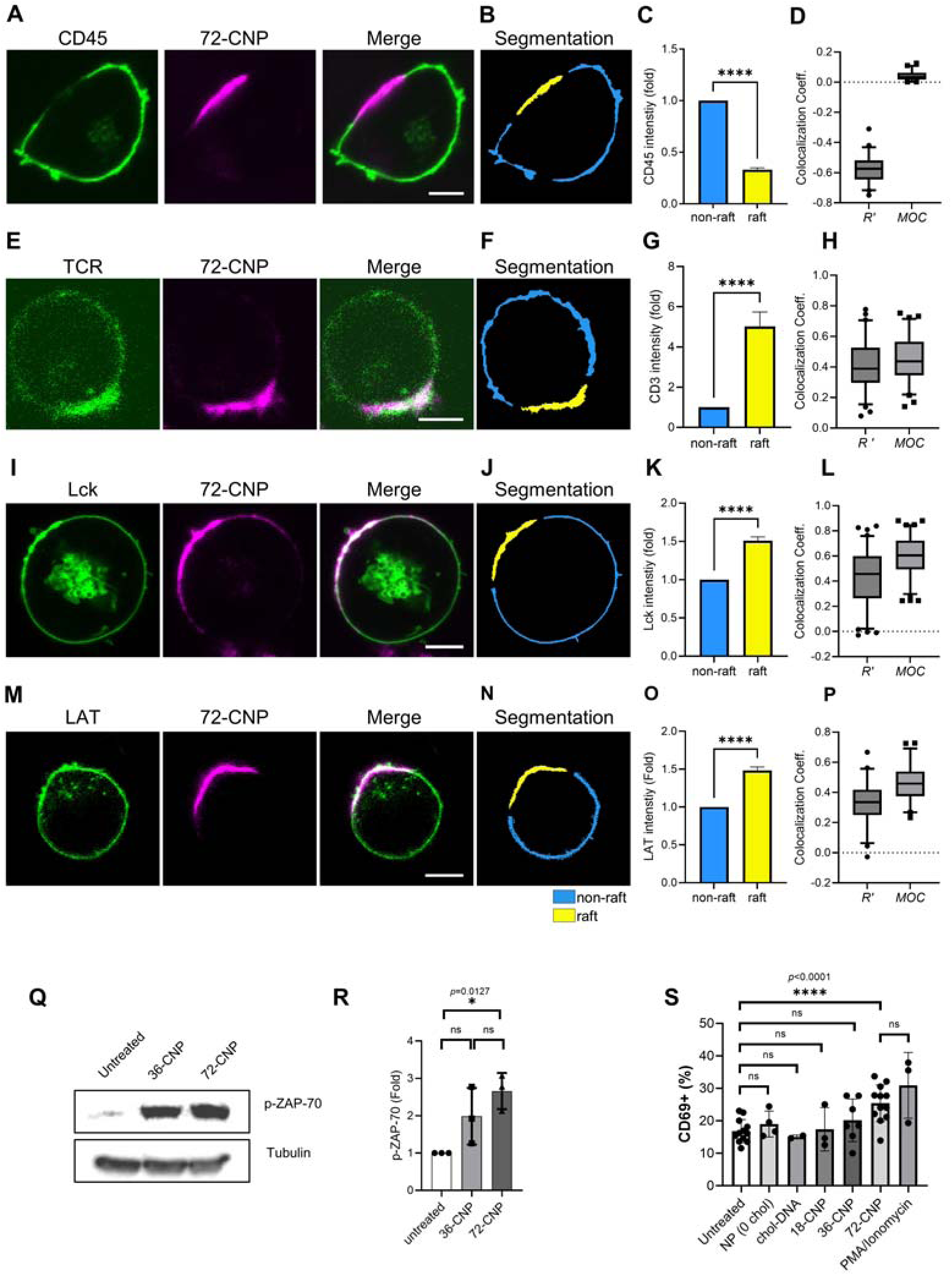
Reorganization of key membrane proteins at the synthetic lipid raft induces early T cell activation. **(A-D)** CD45 is excluded from the synthetic lipid raft. Representative confocal images of CD45 (green, left), 72-DNA (magenta, middle), and the merged image (right) of a Jurkat T cell incubated with 72-CNP for 30 min (A). Segmented regions of the raft (yellow) and non-raft (light blue) regions of the cell in (A) (B). The means of the median intensities of CD45 (normalized) within the segmented non-raft and raft regions of the cells were calculated (n=51 cells collected from three independent experiments) (C). Colocalization coefficients (Pearson’s *R*’ and *MOC*) between CD45 and 72-CNP within the segmented regions of the data set in (C) were calculated (D). (**E-P)** TCR (CD3) (n=68 cells) (E-H), Lck-GFP (n=86 cells) (I-L), and LAT-GFP (n=47 cells) (M-P) accumulations at the synthetic lipid raft, analyzed as described in (A-D), respectively. All data were collected from at least three independent experiments. **(Q-R)** Immunoblot analysis of phosphorylated ZAP-70 (p-ZAP-70) in Jurkat T cells incubated without or with 36- or 72-CNPs for 30 min. Tubulin levels represent the loading control. A representative image of immunoblotting (Q) and quantification of p-ZAP-70 levels (R). The intensity of each p-ZAP-70 band relative to respective tubulin or β-actin band was normalized to that of untreated samples. Each dot represents the band intensity quantified from an individual immunoblotting from three independent experiments. **(S)** Flow cytometry analysis of CD69 expression induced by CNPs in Jurkat T cells. The increases in the CD69+ population of cells in untreated Jurkat T cells, and those treated with NPs without cholesterol (0 chol), 720 nM of Chol-DNAs (equivalent to 10 nM of 72-CNP), and 18-, 36-, and 72-CNP, or PMA/Ionomycin after 4 hours of incubation were calculated. Each dot represents a biologically independent sample encompassing 10,000 cells, derived from at least three independent experiments, except for the Chol-DNA-treated data, which were derived from two independent experiments. Data in (C), (G), (K), and (O) represent means ± SE; Data in (R) and (S) represent means ± SD. *p*-values (*, *p* ≤ 0.05; ****, *p* ≤ 0.0001) were calculated using a two-tailed ratio paired t-test. In (D), (H), (L), and (P), the lines within the boxes represent the means, while the dots above and below the whiskers represent outliers that fall outside the 95th or 5th percentile. Scale bars in (A), (E), (I), and (M): 5µm.

We next investigated whether the synthetic lipid rafts could recruit TCRs, similar to TCR clustering at the core of the immunological synapse ^48^. In 81% of CNP-polarized cells (n=515), we observed CD3 enrichment at the CNP-polarized synthetic raft regions (**Fig. 5E-H**). The average of the median intensities of CD3 within the segmented raft regions was five times higher than that of CD3 in non-raft regions in 68 cells (**Fig. 5F,G**). These results indicate that synthetic lipid rafts drive the segregation of CD45 from TCR, with CD45 excluded and TCR recruited to the rafts.

In addition, we examined the localizations of lymphocyte-specific protein tyrosine kinase (Lck) and linker for activation of T cells (LAT) proteins at the plasma membrane. Lck is one of the earliest signaling molecules that phosphorylate immunoreceptor tyrosine-based activation motifs (ITAMs) on TCR, and LAT is an adaptor protein involved in early TCR signal transduction. Both Lck and LAT are known to be recruited to lipid rafts during T cell activation via myristoylation or palmitoylation ^49–52^. To observe the recruitment of these proteins to the synthetic lipid raft, we utilized Jurkat T cells stably expressing either GFP-tagged-N-terminus part of Lck including the lipidation sites (Lck-GFP) or GFP-tagged LAT (LAT-GFP). We observed the accumulation of both Lck-GFP (**Fig. 5I-L**) and LAT-GFP (**Fig. 5M-P**) within the CNP-induced synthetic lipid rafts. The average of the Lck-GFP and LAT-GFP median intensities was 1.51 and 1.48 times higher in the 72-CNP-induced raft regions than in the non-raft regions in the 86 and 47 analyzed cells, respectively (**Fig. 5J,K**; **Fig. 5N,O**). These results demonstrate that CNP-induced synthetic lipid rafts successfully mimic the function of native lipid rafts at the immunological synapse that recruit early signaling proteins.

We then examined the phosphorylation level of the cytosolic kinase TCR-zeta chain-associated 70-kDa tyrosine phosphoprotein (p-ZAP-70), one of the early downstream events of TCR signaling ^53,54^. While 36-CNP treated cells showed only a modest enhancement of p-ZAP-70 (2.0 ± 0.76 folds), 72-CNP treatment induced a significant increase in p-ZAP-70 (2.7 ± 0.5 folds) compared to the untreated Jurkat T cells (**Fig. 5Q,R**). In human primary effector T cells, we also could observe 1.76 and 2.19 folds higher level of p-ZAP-70 in 36-CNP- or 72-CNP-treated cells, respectively (**Fig. S8A,B**).

We also assessed the surface expression of CD69, an early cellular marker of T cell activation ^55^. Jurkat T cells were treated with or without CNPs and then cultured in complete media for 4 hours before analyzing CD69 expression using flow cytometry. The population of the CD69^+^ cells significantly increased in the presence of 72-CNPs compared to untreated controls (**Fig. 5S**). The level of CD69 increment in the 72-CNP treated samples was not significantly different from that of the PMA/ionomycin-treated samples. However, cells treated with 18-CNP and 36-CNP did not show significant upregulation of CD69. Treating cells with either NPs without cholesterol (0 chol) or 720nM Chol-DNAs (equivalent amount to 10nM of 72-CNP, without origami assembly) alone also did not alter CD69 expression. (**Fig. 5S**).

In summary, these results demonstrate that the creation of cholesterol-mediated lipid rafts is sufficient to induce rearranging the spatial distribution of TCR, Lck, LAT, and CD45, thereby transmitting early T cell activation signals.

## Discussion

In this study, we demonstrated a novel strategy by applying DNA nanotechnology to create synthetic lipid rafts on live T cell membranes that effectively drive lipid phase separation, membrane protein segregations and early T cell activation. Previous studies with model membranes have shown that high cholesterol levels can induce a large-scale LLPS of the lipids ^14,15^. However, such cholesterol-induced membrane segregation on a micrometer scale has not been observed in live cell membranes. The manipulation of phase separation in biomolecules using DNA nanotechnology has gained significant attention in recent years ^56^. DNA, being a programmable and versatile molecule, can be engineered to induce and regulate the phase separation of biomolecules. In this study, we utilized DNA nanotechnology to induce large-scale phase separation of live T cell membranes. The DNA-based CNP platform allows precise and facile control over the number, distance, or arrangement of cholesterols on one side of the DNA origami surface, mimicking a nano-sized lipid raft monomer. The binding of CNPs induced a large-scale LLPS of the membrane, leading to the formation of synthetic lipid raft. The array of multiple cholesterol molecules on the CNPs facilitated their robust and stable binding to T cell membrane, increasing the local cholesterol concentration (**Fig. 6**). CNPs with higher numbers of cholesterols were more effective in driving the segregation of membrane domains, suggesting that changes in local concentration of cholesterol shift the thermodynamic equilibrium, promoting large phase separation in the live cell membrane. This synthetic lipid raft colocalizes with the native lipid raft protein Flot1 and recruits TCR, Lck, and LAT while excluding CD45, thereby inducing early T cell activation (**Fig. 6**).

**Figure 6.**
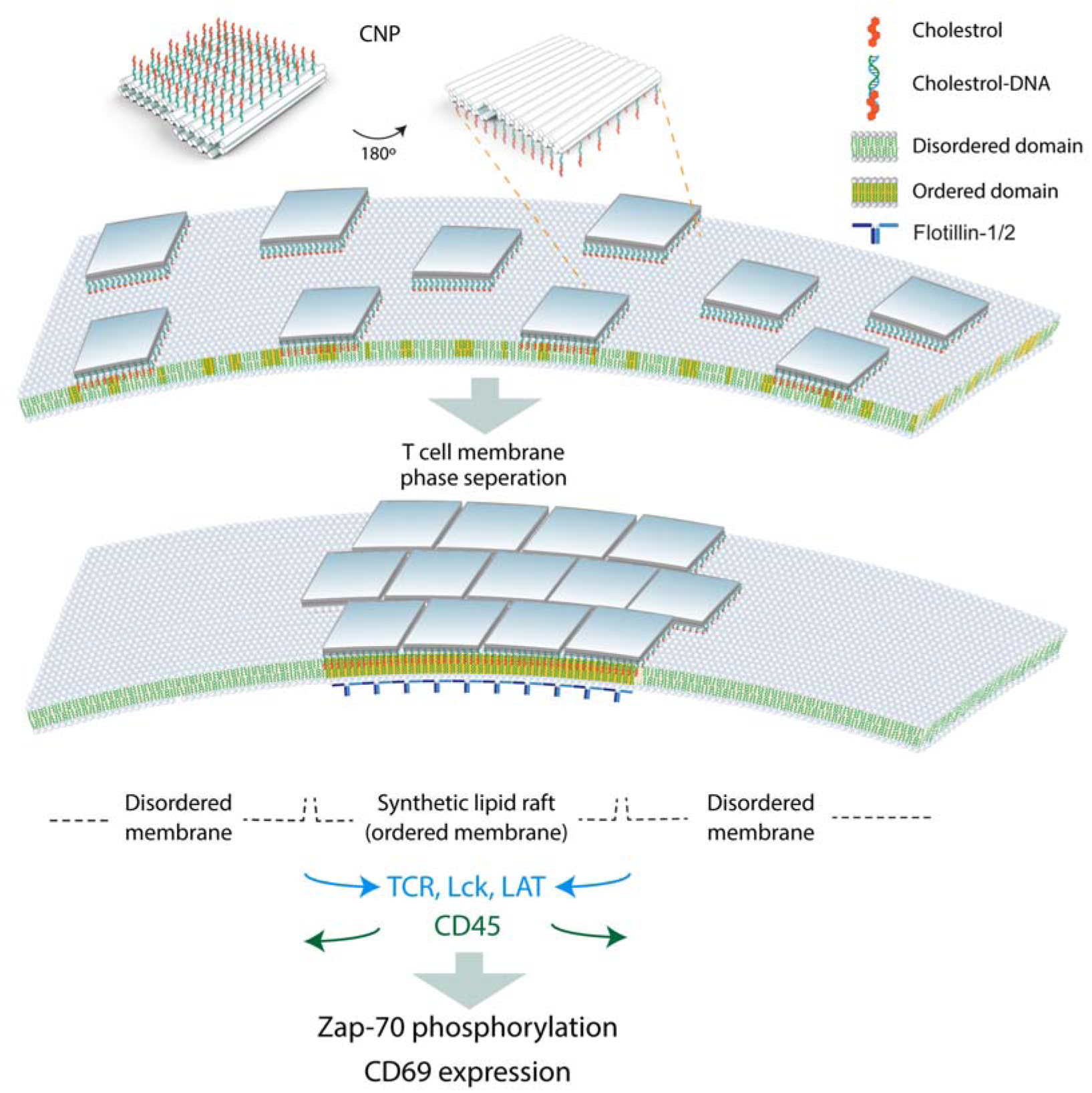
Proposed model of CNP-mediated synthetic lipid raft formation on the T cell membrane and its effects on early T cell activation. Binding of 72-CNPs induces phase separation of lipid domains in the T cell plasma membrane, resulting in the formation of synthetic lipid rafts. These rafts recruit TCR, Lck, and LAT while excluding CD45, thereby initiating ZAP-70 phosphorylation and CD69 expression—hallmarks of early T cell activation.

Our study directly demonstrates the role of cholesterol in the formation of lipid rafts on the live T cell membrane and their involvement in the reorganization of key membrane proteins. Although the spatial redistribution of these receptors has been well-documented, it remains unclear whether lipid rafts drive this rearrangement or whether lipid raft coalescence is a consequence of receptor reorganization. Our results indicate that the formation of lipid rafts is sufficient to reorganize membrane proteins according to their preference for these domains. Our results showed that ∼ 67 % of CD45 molecules were excluded from the synthetic lipid rafts in the absence of forming close contact with an APC. This result supports the mechanism of lipid raft-driven CD45 segregation and is aligns with hypothesis that cholesterol can cause the segregation of CD45 ^47^. The incomplete segregation of CD45 in our study, along with previous reports from reconstitution studies involving modifications in ligation length ^57,58^ suggests that CD45 exclusion at the immunological synapse is driven by both lipid raft-mediated and extracellular domain mediated-kinetic segregation mechanisms.

We observed a high colocalization of CNPs with the preassembled Flot1 cap on T cells. Previous studies have reported that Flot1 preassembled cap is resistant to cholesterol-disrupting agents ^38^. Our results show that the assembly of Flot1 could be initiated and enhanced by increasing the local concentration of cholesterol. Additionally, we assume that the cholesterol molecules complexed with the CNP insert exclusively into the outer leaflet of the cell membrane because the bulky DNA origami structure is unlikely to undergo flip-flop. The significantly decreased diffusion of Flot1 upon the CNP binding indicates that the concentration of cholesterol in the outer leaflet can induce the transbilayer coupling with the adjacent inner leaflet of the plasma membrane ^59^.

The biological significance of lipid rafts has been a controversial topic for decades. Our strategy for controlling the spatial distribution of cholesterol can be easily modified or extended to include other major components of lipid rafts, such as sphingolipids or other proteins, in various cell types. Our DNA nanostructures may have ramifications for the design of novel molecular therapeutics capable of controlling T cell immune responses independent of specific antigens, thus expanding the potential of T cell-based immunotherapy. The CNP can be applied to manipulate the phase separation of various biomolecules to modulate cellular processes. Furthermore, our cholesterol patterning technology can be combined with arrangement of specific antigen peptides on DNA origami that might bring synergic effects on activating antigen-specific T cells or can be utilized for the design of CAR-T cell with enhanced therapeutic effects.

## Materials and Methods

### Mice

C56BL/6J mice (8-18-week-old male) were purchased from Orient Bio. Mice were maintained and used by following the guidelines of the Yonsei Univesity Institutional Animal Care and Use Committee (IACUC), and approval for use of mice was obtained from Yonsei Univesity IACUC (ACUC-A-202306-1687-01(M.K.))

### Cells

Jurkat T cells (Clone E6-1, ATCC) and Jurkat T cells transduced with Flot1-Halo, Lck-GFP, or LAT-GFP were cultured in complete RPMI (Gibco) media containing 2 mM L-glutamine, 100 U penicillin-streptomycin (PS) (Gibco) and 10% FBS (Gibco). Human pan T cells were isolated from frozen peripheral blood mononuclear cells (PBMCs, STEMCELL Technologies, Cat #70025.2. Lot 201671701C) using the negative selection human Pan T Cell Isolation Kit (Miltenyi Biotec). Mouse T cells were purified from mouse splenocytes using the negative selection EasySep™ Mouse T Cell Isolation Kit (STEMCELL Technologies). Mouse splenocytes were optained from 8-18-week-old male BL6 mice purchased from Orient Bio. For each experiment, a spleen was processed through a 40 μm filter (STEMCELL Technologies) and RBCs were removed from the single cell suspension using RBC lysis buffer (eBioscience). Freshly isolated resting cells for both human and mouse were used after incubation in complete T cell media, containing 10% (v/v) FBS, 100 U PS, 1x GlutaMAX supplement (Gibco), 1mM sodium pyruvate (Sigma Aldrich) and 1x nonessential amino acids solution (Sigma Aldrich) in RPMI, for ∼2-4 h in a CO_2_ (5%) incubator at 37 °C. For human effector T cells, purified pan T cells were cultured with human anti-CD3/anti-CD28 Dynabeads (Gibco) at a 2:1 ratio in complete T cell media supplemented with 50μM 2-mercaptoethanol and 100 U·mL^−1^ of IL-2, for up to two weeks.

### Synthesizing the cholesterol nano-patch (CNP)

The DNA nano-patch (NP) was designed on the honeycomb lattice with a long single-stranded scaffold, type p8064 (Tilibit), by modifying the structure of R. Dong et al. ^60^ Sequence of 183 staple strands of the DNA NP was determined using the open-source program, caDNAno,^61^ (**Table S1**) and the corresponding staple strands were synthesized from Bioneer. A folding mixture consists of 20 nM of scaffold DNA, 100 nM of each staple strands, 20 mM MgCl_2_ and 5 mM NaCl in TE buffer (5 mM Tris and 1 mM EDTA, pH 8). The annealing was performed following a temperature profile: 70 °C for 15 min, 60 to 20 °C by 1 °C/ho in a thermocycler. Excessive staple strands were removing by polyethylene glycol (PEG) precipitation ^62^ followed by two rounds of ultrafiltration with a 100 kDa cut-off filter. A concentration of the DNA NP was measured using UV spectrometer and adjusted using the working buffer (1×TE, 10 mM of MgCl_2_, and 5 mM of NaCl). Purified samples were stored at 4 °C. To form the CNP, we mixed the DNA NP with four-fold excessive assist-DNAs and Chol-DNAs to the total number of DNA handles on the DNA NP. The mixture was then incubated them at 30 °C for 12 h. At the 3’ end of assist-DNAs, a six-nucleotides-thymine (6T) is appended to minimize undesired hydrophobic interaction between Chol-DNAs by wrapping the neighboring cholesterol ^63^. To remove undesired hydrophobic interactions, Chol-DNA was heated at 65 °C for 1 h before use. The mixed solution was purified by ultrafiltration with a 100 kDa cut-off filter using phenol red free RPMI (Gibco) and its concentration was adjusted using the phenol red free RPMI. To evaluate the formation of complexes with the desired number of CNPs on the specific available binding sites on NPs, we replaced cholesterol with Cy5 fluorophore in Chol-DNA and measured the fluorescence intensities of the complexes at 670 nm with 649 nm excitation using a fluorometer (FluoroMax Plus, Horiba) and a FluorEssence (version 3.8, Horiba) software.

### Agarose gel electrophoresis

The assembled NP was electrophoresed using 1% agarose gel containing 0.5×TBE (45 mM Tris-borate and 1 mM EDTA), 10 mM of MgCl_2_ and 0.1 ul·ml^−1^ of SYBR Safe (Thermofisher). Electrophoresis was performed for 90 min at 70 V bias voltage in a cold room. Agarose gel was scanned using a GelDoc and Image Lab program (version 5.1, Bio-Rad).

### TEM Imaging

The concentration of CNP was diluted to 3 nM in the working buffer. 5 μL sample solution was deposited on a glow-discharged formvar-supported carbon-coated grid for 3 min and stained by a 2% uranyl formate and 25 mM NaOH for 40 sec. TEM images were taken at 40k-fold or 80-fold magnification using JEM-2100Plus (JEOL).

### Sample preparation for imaging

Jurkat or primary T cells were washed with calcium and magnesium free PBS (Gibco) through centrifugation at 200 g for 5 min (spin wash). The cells were then resuspended in phenol red free RPMI. For the sequential reaction, cells were spin washed and subsequently incubated with 10 µM of Chol-DNAs for 10 min at 37 °C. After another spin wash, 10 µM of DNA assist-DNAs was added and incubated for 10 min. Following this, 10 nM of DNA NP (containing 72-anchors) was added to the sample. The unbound DNAs were eliminated through a spin wash. For the single step reaction, Chol-DNAs were pre-complexed with DNA NPs prior to use. The CNPs were then directly added to the spin washed cells at a final concentration of 10 nM in a total volume of 50 µl of phenol red free RPMI and incubated in a CO_2_ (5%) incubator for 30 min at 37 °C, or, as indicated. Subsequently, the cells were spin washed with PBS or phenol red free RPMI, ready for using live cell imaging. For nuclei staining, cells were fixed with 4 % (w/v) paraformaldehyde (PFA) (Electron Microscopy Sciences) in PBS for 15 min at room temperature (RT), and subsequently washed in PBS. Cells were stained with 10 uM of Hoechst 33342 (Thermo Fisher Scientific) for 5 min, followed by a wash with PBS. For Flot1-Halo labeling, after washing cells with PBS, they were stained with Janelia Fluor (JF) 549 conjugated Halo-tag ligands (Promega) at 200 nM concentration in 2% FBS/PBS at 37 °C for 15 min. The unbound Halo-tags were removed through a spin washed with PBS. The cells were then incubated with CNPs for 30 min, as described above. For CD45 labeling, the CNP labeled cells were washed and blocked with 2% BSA in PBS for 10 min in a poly L-lysine (PLL) coated glass bottom chamber. They were then labeled with 1ug·ml^−1^ of Alexa Flour (AF) 488 conjugated anti-human CD45 antibodies (Biolegend, Cat# 304017, clone HI30) in 2% BSA in PBS for 15 min at RT. After labeling, cells were rinsed with PBS four times. For CD3 labeling, the CNP labeled cells were spin washed and incubated with 10µg·ml^−1^ of AF488 conjugated anti-CD3 antibodies (Biolegend, Cat # 300415, clone UCHT1) in 2% BSA/PBS on ice for 30 min. After incubation, the cells were spin washed again. For imaging, 70∼100 µl of freshly prepared cells resuspended in PBS were added to a PLL (0,01 %, Sigma Aldrich) or collagen (50 mg·ml^−1^, Corning) coated glass-bottom chamber (35 mm dish, 7 mm glass, MatTek). The chambers were freshly prepared prior to use. For live time-lapse imaging, Halo-tag labeled cells resuspended in phenol red free RPMI were placed on the PLL coated glass bottom chamber followed by adding 3-5 µl of CNP in RPMI to achieve a final concentration of 3-5 nM. Image acquisition was initiated 2 min after adding the CNPs.

### Plasmid construction and lentiviral transduction

The lentiviral pLV-FLOT1-Halo plasmid was gifted from Prof. Young-Wook Jun (University of California San Francisco). To construct the lentiviral pLV-Lck-GFP plasmid, partial Lck (135 bp)-EGFP sequences were amplified from pBOB-Lck-GFP (Addgene # 118738) using PCR with the following primers: forward: 5’-cgggcctttcgaattccgctgttttgacctccat-3’ and reverse: 5’- gtgggagttgcggccgctcagcattggtagctgct −3’. The lentiviral vector pHR_PGK_antiCD19_synNotch_Gal4VP64 (Addgene # 79125) was digested with EcoRI and NotI, and then ligated with the amplified sequences. To construct the lentiviral pLV-LAT-GFP plasmid, the LAT sequence was amplified from the pHR LAT/Zap70 iLID-Drop plasmid (Addgene # 171038) by PCR using primers incorporating the EcoRI enzyme restriction sites: forward: 5’-tatttccggtgaattatggaggaggccatcctggtc-3’ and reverse: 5’- tcaccatggtgaattcgttcagctcctgcag-3’. The amplified sequences were then cloned into the lentiviral vector pLVX-EF1alpha-eGFP-2xStrep-IRES-Puro (Addgene #288041) at the EcoRI enzyme restriction site (before GFP).The resulting vectors were confirmed by DNA sequencing. The viral plasmid and two packaging plasmids pSPAX2 (Addgene) and pMD2.G (Addgene) were cotransfected into Lenti-X293T cells (Takara) using lipofectamine 2000 or lipofectamine 3000 (Thermo Fisher Scientific) following the manufacturer’s instructions. After 48 hours, the supernatant was collected and filtered using a 0.45 µm pore sized Millex PVDF syringe filter (Millipore). The filtered virus was concentrated by ultracentrifugation at 10,000 g for 2 h and then applied to Jurkat T cells for a two-day transduction period. Following transduction, cells were subjected to spin washing and were maintained in complete RPMI media for three days. Afterwards, cells were harvested for sorting. The sorted cells were cultured in complete T cell media for additional two days and then maintained in complete RPMI media.

### Flow cytometry

The bindings of DNA NP, with or without cholesterol modification, to single cell were analyzed using flow cytometry. After incubating the cells with DNA NP or CNP for 30 min, they were either washed or supplemented with 1 ml of complete RPMI media, following by a 4 h incubation at 37 °C in a CO_2_ (5%) incubator. The cells harvested after 30 min or 4 h incubation were spin washed with PBS and then stained with LIVE/DEAD™ Fixable Violet Dead Cell Stain dye (L/D violet) (Invitrogen) diluted in PBS (1:2000) for 10 min at RT in the dark. Subsequently, the cells were spin washed and loaded on a Fusion cell sorter (BD Biosciences) for analysis. For flow cytometric analysis, single cells were first gated using forward vs. side scatter (FSC vs. SSC), and then live cells were gated based on the negative population of the L/D violet stain. The Cy5 fluorescence on CNPs was analyzed using FlowJo (BD Biosciences, versions 10.8.1) software. For the analysis of CD69 expression, cells were incubated in RPMI without CNP, with CNP, or with PMA (10 ng·ml^−1^) and ionomycin (0.5 µg·ml^−1^) (both from Sigma Aldrich) at 37 °C for 30 min. Then, 1 ml of complete RPMI was added, and the cells were further incubated at 37 °C for 4 h in a CO_2_ (5 %) incubator. Subsequently, the cells were harvested and spin washed with PBS. The cells were then labeled with L/D violet for 10 min at RT. After another spin wash with 2% FBS/PBS, the cells were resuspended in 100 µl of 2 % FBS/PBS and stained with Brilliant Violet (BV) 785 conjugated anti-human CD69 (10 µg·ml^−1^) (Biolegend, Cat # 310932, clone FN50) for 30 min on ice. Cells were spin washed and then loaded on a FACSAria Fusion cytometer (BD Biosciences). For compensation, ArC amine reactive compensation beads (Invitrogen) stained with L/D violet and UltraComp eBeads (Invitrogen) stained with AF 647 conjugated anti-IgG antibody for Cy5 or the BV785 conjugated CD69 antibody in PBS were used. The live single cell population positive for CD69 was analyzed using the FlowJo software. For cell proliferation tracking analysis, Jurkat T cells were harvested and spin washed in PBS. They were then stained with Cell Tracker Green CMFDA (1 µM) (Invitrogen) diluted in 2% FBS/PBS for 20 min at 37 °C in a CO_2_ (5 %) incubator. After spin washing with 2 % FBS/PBS, cells were resuspended in RPMI and incubated without or with 36- or 72-CNP at a concentration of 10 nM for 30 min at 37 °C in a CO_2_ (5 %) incubator. The cells were spin washed (day 0 sample) or grown in complete RPMI media for 5 days. Cells were harvested and the diluted intensities of the CMFDA were analyzed every day (day 0–day 4) using flow cytometry. After spin washing the harvested cells, they were stained with L/D violet in PBS (1:2000) for 10 min at RT. Following this, cells were rinsed with PBS and loaded onto the cytometer. The distribution of CMFDA intensities on the live Jurkat T cells was analyzed using the FlowJo software.

### Fluorescence-activated single cell sorting (FACS)

For sorting, Jurkat T cells transduced with Flot1-Halo, Lck-GFP, or LAT-GFP were spin washed with PBS and stained with L/D violet (1:2000) for 10 min at RT. Flot1-Halo transduced cells were spin-washed again and stained with JF549-conjugated Halo-tag ligands (Promega) at a concentration of 200 nM in 2% FBS/PBS for 20 min at RT. The cells were underwent another spin wash with 2 % FBS/PBS. The cells underwent another spin wash with 2% FBS/PBS, were resuspended in 2% FBS/PBS, and kept on ice in the dark. Immediately afterward, cells were loaded onto a Fusion cell sorter (BD Bioscience). Live single cells were gated as previously described, and the JF549 or GFP positive cells were analyzed and sorted using FACSDiva software (BD Biosciences, version 8.0.2).

### Confocal imaging

Confocal images were acquired using two different microscopes: a laser scanning Ti Microscope (Nikon) equipped with an Apo 60 x Oil λS DIC N2 objective lens (N.A., 1.4) (Nikon), and a Zeiss LSM 900 (Zeiss) microscope, equipped with a Plan-Apochromat 63x Oil DIC M27 (N.A. 1.4) (Zeiss) objective lens. The pinhole size used ranged from 46 to 52 µm. For the Nikon microscope, four laser lines at 405, 488, 561, and 633 nm were employed, along with a combination of filter sets (dichroic: BS 20/80, emission bandpass: 450/50, 525/50, 595/50, and 700/75). Bright field images were taken with a transmission detector. For the Zeiss microscope, 405, 488, 561, and 640nm excitation laser lines were used, and detection spectra for each fluorophore were determined using two variable beam splitter dichroics (VSDs).

### C-laurdan imaging

CNP-labeled cells were stained with 10 µM of C-laurdan dye for 10 min at 37 °C in a CO_2_ (5 %) incubator. Confocal images were acquired using a Zeiss LSM 900 (Zeiss) microscope, equipped with a Plan-Apochromat 63x Oil DIC M27 (N.A. 1.4) (Zeiss) objective lens with a 50 µm sized pinhole. For excitation, the C-laurdan and Cy5-labeled CNP were illuminated using laser lines at 405 nm and 633 nm, respectively. Emission fluorescence within the ranges of 400-460 nm and 470-530 nm was collected to calculate the GP value as described previously ^64^. The GP value is calculated as: 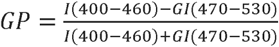, where I(400-460) and I(470-530) are the fluorescence intensities of each pixel in the image collected within the 400-460 nm and 470-530 nm, respectively and G=(GP_ref_+GP_ref_*GP_mes_-GP_mes_-1)/ (GP_mes_+GP_ref_*GP_mes_-GP_ref_-1). Here, GP_mes_ is 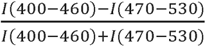, measured with C-laurdan dye (10µM in DMSO), and GP_ref_ is the reference value that makes the mean GP of the plasma membrane becomes 0. To segment the CNP-bound (raft) regions, CNP emission between 645-700 nm was obtained simultaneously with each channel acquisition.

### FRAP analysis

For FRAP experiments, time lapse confocal images were acquired using a Zeiss LSM 900 (Zeiss) microscope, equipped with a C-Apochromat 40x Water Korr (N.A. 1.2) (Zeiss) objective lens with a 50 µm sized pinhole. Laser lines at 561 nm and 633 nm were used for excitation, and bleaching and emission fluorescence between 400-635 nm and 645-700 nm were collected for the JF549-Flot1-Halo and Cy5-labeled CNP, respectively. Time-lapse confocal images were taken over a 5 min period with intervals of either 3 s or 10 s. For the photobleaching step, after taking 3-5 frame images, a squared area of 0.5 µm^2^ was selected, and this area was bleached for 3 s using 80 % power for each laser line. To correct for any drifting or motility of cells during the measurement, each frame image was adjusted to align with the next frame image, except for the bleached frame. This alignment was performed using the ‘imregtform’ function in MATLAB (MathWorks, MathWorks, version R2018a, version R2023a), with the ‘rigid’ optimizer option. The mean fluorescence intensities of the bleached area over the course of the time-lapse movie were tracked using a custom MATLAB script. The diffusion coefficient (*D*) for Flot1 was determined by analyzing the fluorescence recovery trace during the 2.5 min following bleaching. The analysis was conducted using the following equation ^65^: 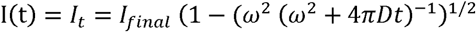, where *I*_*t*_ is intensity as a function of time, *I_final_* is final intensity reached after recovery, *ω* is width of the bleached area, and *D* is one dimensional diffusion coefficient, which was determined through fitting using a built-in function in MATLAB. For FRAP analysis, images that have significant and uncorrectable sample drifting during the image acquisition were excluded from the data analysis.

### Image analysis

ImageJ (version 1.52p) and MATLAB were used for image processing. For counting the polarization population, each population of cells was counted from the maximum intensity projection of 9-11 z-stack images using an ImageJ built in function, cell counter. The perimeters of the single polarized clusters were manually measured from a z plane of the middle of each cell in the confocal images using image J. For the colocalization test between channel X and channel Y, Pearson’s correlation coefficient ^66^ 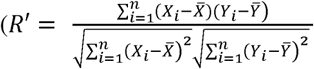, where *n* is the total number of pixels, *X_i_* and *Y_i_* are the pixel intensities indexed with *i*. 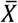 and 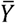 is the mean intensity of *X* and *Y* images, respectively) and Manders’ overlap coefficient ^67^ 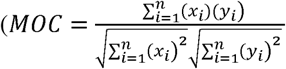, where *n* is the total number of pixels, *x_i_* and *y_i_* are the intensities of above-threshold pixels indexed *i* in the *X* and *Y* images) were calculated using a custom built MATLAB script. The original intensity values of the membrane regions were utilized without subtracting background when calculating *R’*. For *MOC,* the initial threshold values were determined using the ‘multithresh’ function in MATLAB. To quantify the fold differences in Flot1-Halo, CD45, CD3, Lck-GFP, and LAT-GFP intensities between the raft and non-raft regions, median intensity for pixels within each single cell’s raft and non-raft areas were normalized by the median non-raft intensity. Background values, represented by the minimum intensity for each image, were subtracted during the intensity analysis.

The region of interest (ROI) for individual cells was selected using ImageJ or MATLAB script. To segment the CNP-bound (raft) and unbound (non-raft) regions of the plasma membrane of the dual-labeled images, a custom MATLAB script was employed. Briefly, channel 1 (Flot1-Halo, CD45, CD3, Lck-GFP, and LAT-GFP) and channel 2 (72-CNP) images, along with the summed images of these two channels, were thresholded using the threshold value obtained from the ‘multithresh’ or ‘imbinarize’ function in MATLAB. This created three binary images: BW1 (for channel 1), BW2 (for channel 2), and BW3 (for both). Noise elements consisting of fewer than 5 connected components were removed from BW3 using the ‘bwconncomp’ function. To bridge the gaps between the disconnected membranes, BW3 was dilated using a spherical structuring element with a radius corresponding to the estimated gap size. Subsequently, a skeleton binary image was generated using the ‘bwskel’ function to dilate the skeleton of the previously dilated BW3, employing a spherical structuring element with a radius of 1. The connected membrane binary image (MB) was obtained by combining the dilated BW3 and the skeleton binary image. For low-signal images causing rough edges on MB, smooth the binary image using the ‘conv2’ function with a 5×5 uniform kernel convolution, followed by rethresholding to generate a binary output. Next, morphological closing was applied to the nonzero elements of BW2 within the binary image MB to define the ‘raft’ binary image, and holes in the binary image were filled using the ‘imfill’ function. The rest of the raft region within the binary image define the ‘non-raft’ binary image. Any remaining elements with less than 20 or 50 connected components were eliminated from the segmented raft or non-raft areas using the ‘bwconncomp’ function, respectively. The boundary regions (1µm) between the raft vs non-raft were eliminated. The nonzero element of raft or non-raft became the total membrane segmented area.

To segment the raft and non-raft regions of the plasma membrane in C-laurdan imaging, the summed image of the two C-laurdan emission channels and the CNP images were used. The ‘imbinarize’ function with ‘adaptive’ option (sensitivity =0.5) was applied to create an initial segmentation of the label cells. To exclude regions other than the plasma membrane, the ‘regionprops’ and ‘bwconncomp’ functions were employed to identify connected components based on attributes such as ‘circularity,’ ‘solidity,’ ‘pixel area,’ or ‘size of the largest hole’ in the threshold binary images. The CNP image was binarized using the ‘imbinarize’ function with the ‘global’ option. Overlapping regions between the segmented plasma membrane and the CNP binary image were identified as raft regions. The remaining portion of the plasma membrane represents the non-raft regions. Objects with fewer than 50 connected components in raft regions and 1000 connected components in non-raft regions were eliminated. The boundary regions (1µm) between the raft vs non-raft were eliminated. The cells and image field of view were selected randomly for each group of experiments. Blinding was performed during image segmentation and analysis. Image analysis was performed automatically, except manual counting for the polarized populations.

The intensity display ranges for individual color channels in **Fig. 2, Fig. 4G, Fig. S2, Fig. S3, and Fig. S5** were independently optimized using ImageJ or MATLAB. However, for comparison purposes, the display range of Cy5 intensity in **Fig. S2B** was adjusted to that of **Fig. 2A**; the 0-, 18- and 36-CNP images in **Fig. 2B and Fig. S3** were adjusted to match the range of the 72-CNP image in **Fig. 2B**; **Fig. S5A** was adjusted to the **Fig. 2E**. The display ranges of the all figures in **Fig. S6** and **Movies S1-S3** were equal. All other images in this study were displayed between the 1% ∼ 99% intensity of individual channel image using a built-in function in MATLAB.

### Immunoblotting assay

Jurkat T cells and primary human T cells were incubated with 10 nM of 72-CNP in RPMI for 30 min at 37 □. The cells were subsequently spin washed with ice-cold PBS and then lysed in lysis buffer containing 20 mM HEPES (pH 7.0), 1 % Triton X-100, 150 mM NaCl, 10 % glycerol, 1 mM EDTA, 2 mM EGTA, 1x protease Inhibitor cocktail (Cell Signaling Technology) and 1x phosphatase inhibitor cocktail (Cell Signaling Technology). The cell lysates were centrifuged at 12,000 g for 10 min, and the protein concentrations were determined by Bradford assay (Bio-Rad) or micro BCA assay (Invitrogen). The protein samples were mixed with SDS sample buffer containing 312.5 mM Tris (pH6.8), 10 % SDS, 30 % glycine, 1mg·ml^−1^ bromophenol blue and 1 M 2-mercaptoethanol, and boiled for 5 min. SDS-PAGE was performed to separate the proteins, which were subsequently transferred to nitrocellulose membranes via electroblotting for 1 h 30 min. Following the transfer, the membranes were blocked with 5 % dry skimmed milk in Tris-buffered saline containing 0.05 % (v/v) Tween-20 (TBST) for 1 h. Subsequently, the membrane were incubated with rabbit anti-phospho ZAP-70(Tyr319)/ Syk (Tyr352) (Cell Signaling Technology, Cat # 2717T, Clone 65E4), anti-β actin antibody (Cell Signaling Technology, Cat # 4970, clone 13E5), or anti-α-tubulin antibody (Sigma Aldrich, Cat #T5168, Clone B-5-1-2) diluted in the blocking buffer (1:1000) for overnight at 4 D. After being washed 3 times with TBST, the membranes were incubated with HRP conjugated anti-rabbit IgG secondary antibody (Cell Signaling Technology, Cat #7074) diluted in blocking buffer (1:2000) for 1 h at RT and the washed with TBST 3 times. The immune-reactive bands were detected using an enhanced chemiluminescence kit (Sigma Aldrich) and ChemiDoc imaging system (Bio-Rad) and quantified with ImageJ software (version 1.52p).

### Statistical information

At least three biologically independent experiments were performed except **Fig. S1, Fig. S2B,** and **Fig. S4** (two independent experiments) and the statistical details for each experiment are indicated in the figure legends. The p-values, mean, standard deviation (SD), and standard error of the mean (SE) were determined using the built-in functions in Matlab (MathWorks, R2018a, R2023a) or Prism 8 and Prism 10 (GraphPad, version 8.0.1, version 10.0.2).

### Data availability

The sequences for the DNA origami structures are listed in the Table S1. Data that are generated and used in this study are available from the corresponding author upon reasonable request.

### Materials availability

Plasmids generated in this study are available from the lead contact. DNA origami made in this study will be made available on request, but we may require a payment and/or a completed materials transfer agreement due to high cost

### Code availability

Custom MATLAB codes used in this study are available upon request to Y.J.

### Reagents and tools table

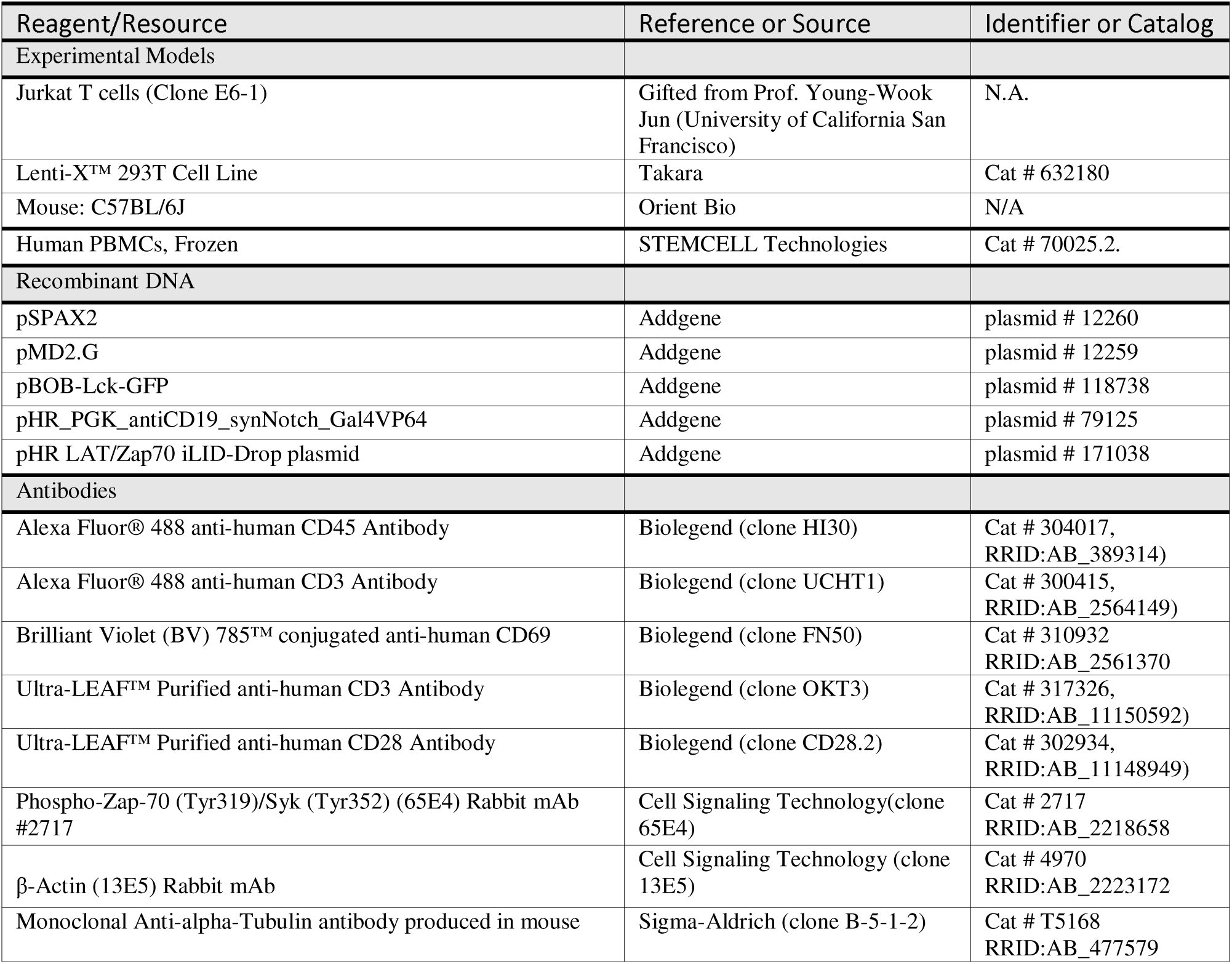
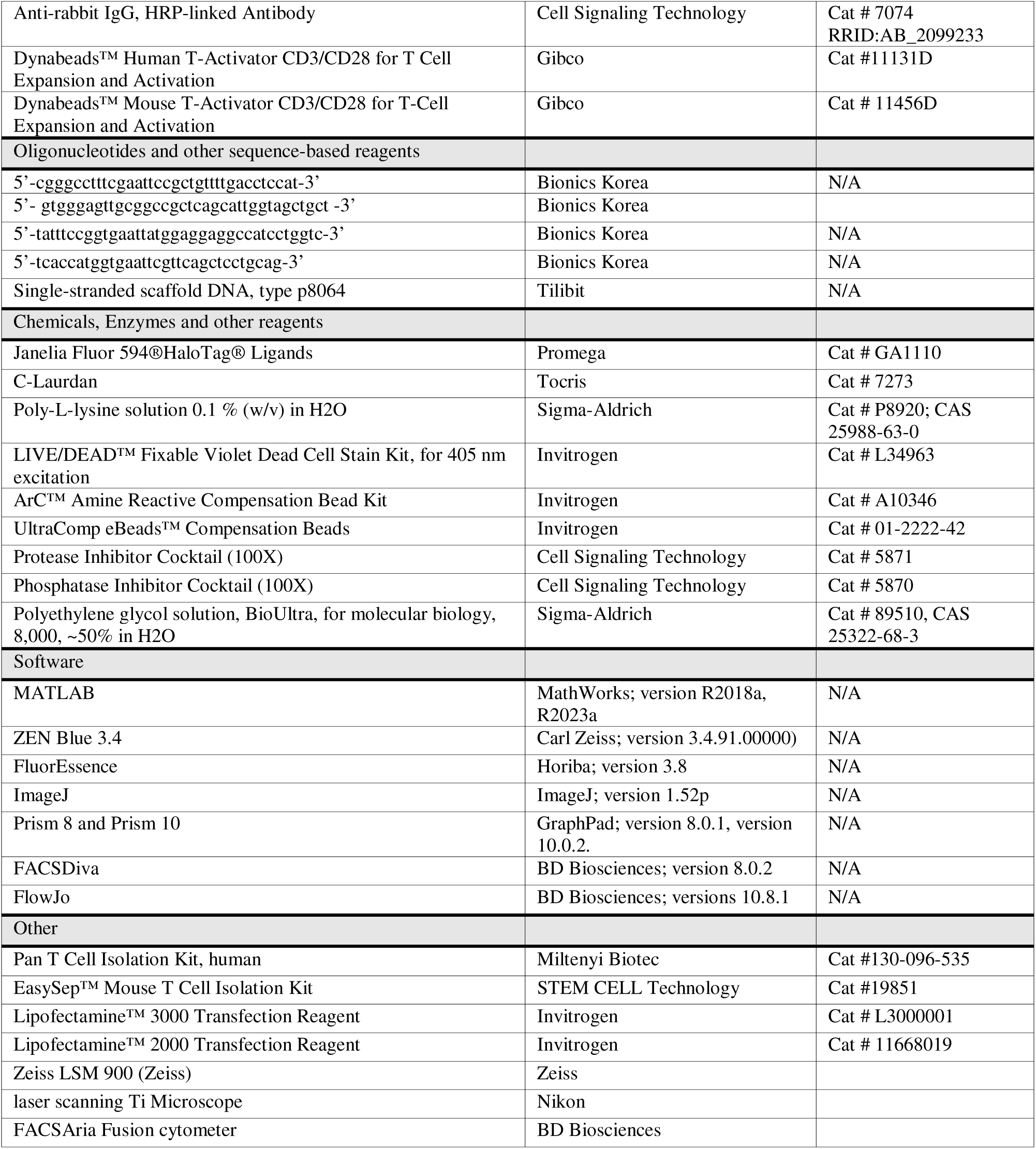

## Acknowledgments

We thank Prof. Gilad Haran (Weizmann Institute of Science), Prof. Ronen Alon (Weizmann Institute of Science), and Prof. Amnon Altman (La Jolla Institute for Immunology) for their kind advice and discussion for the manuscript; Ms. Seonyeong Ha, Jungmin Nam and Mr. Jinyoung Kim for cell culture and technical supports.

## Funding

This research was supported by Basic Science Research Program through the National Research Foundation of Korea (NRF) funded by the Ministry of Education (NRF-2021R1F1A1063378 (M.K.), NRF-2022R1I1A1A01070462 (Y.J.))

## Author Contributions

Y.J. conceived and designed the experiments, Y.J. performed experiments and analyzed data, Y.K. designed the origami construct, Y.K, and K.N synthesized and conducted evaluations for the CNPs. S.L. performed immunoblot assay, Y.J. Y.K. and M.K. wrote manuscript, and M.K. supervised the project.

## Competing Interest Statement

The authors declare no competing interests.

## Notes

### Competing Interest Statement

The authors have declared no competing interest.

### Summary of Updates

Fig 3 is added. In this figure, c-Laurdan staining demonstrates CNP-bound polarized membrane cap-like microdomains stabilize more ordered phase of the plasma membrane. Fig 5I-L and 5M-P for Lck and LAT spatial distribution with respect to CNP-induced synthetic lipid rafts are added. Fig S5 is added showing CNP stimulation can trigger T cell signaling in primary human and mouse T cell.

