## Supplementary Materials for "Synthetic lipid rafts formed by cholesterol nano-patch induce early T cell activation"

**Jung et al.**

**Figure S1 to S8**

**Legends for Movies S1 to S5**

**Tables S1**

**Figure S1**

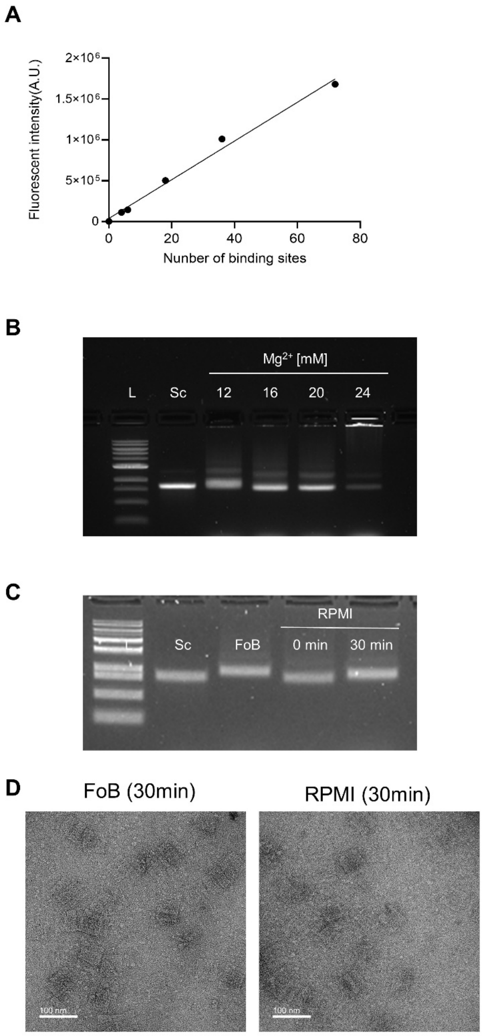


**Figure S1. Folding and assembly of DNA nano-patches (NPs). (A)** Quantification of controlled conjugation. The fluorescence intensities of NPs complexed with DNAs, where cholesterols in Chol-DNAs are substituted with a Cy5 fluorescent dye, exhibited a linear increase dependent on available binding sites. **(B)** Folding of DNA NPs in different magnesium concentrations. Agarose gel electrophoresis confirmed proper folding of DNA NPs in various magnesium concentrations. Well-folded DNA nanostructures were observed in 16 and 20 mM of $Mg^{2+}$ without significant aggregation. In this work, DNA-NP was folded at 20 mM of $Mg^{2+}$, based on these results**. (C-D)** Stable NPs in T cell culture media. Agarose gel electrophoresis (C) and TEM imaging (D) were conducted using purified NPs that were incubated either in the folding buffer or in RPMI for 30 min, the condition used for labeling CNPs to cells in this work. (L: 1kb DNA ladder; Sc: 8064 scaffold strands; FoB: Folding buffer (20 mM $Mg^{2+}$)).

**Figure S2**

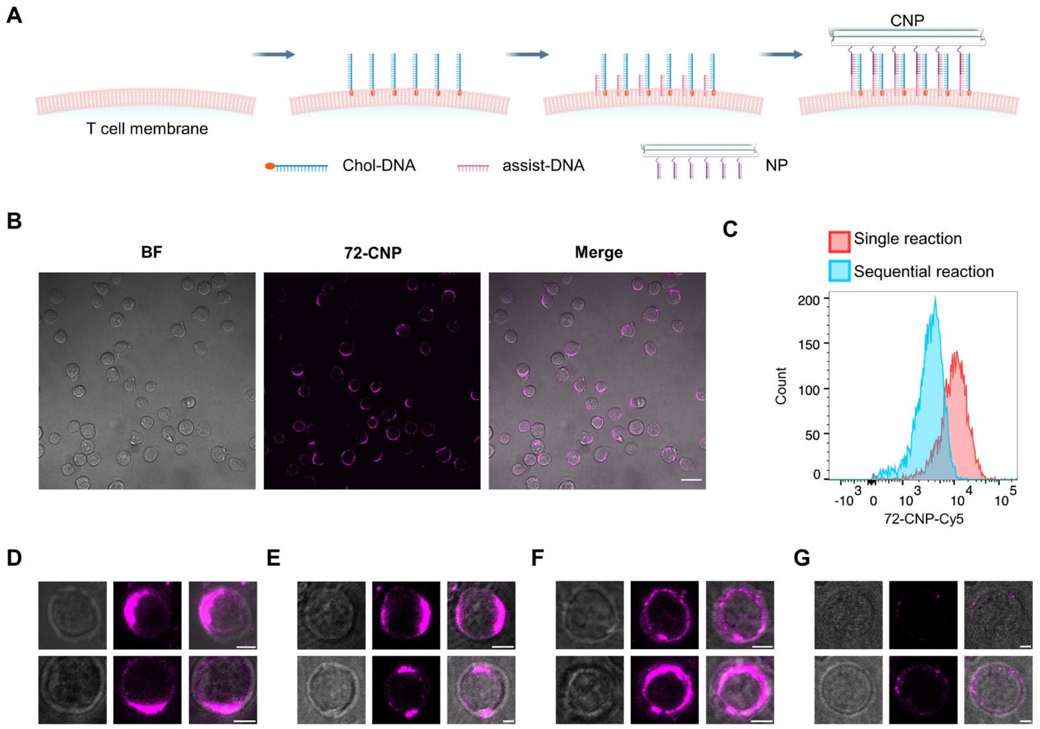


**Figure S2. CNP binding on the plasma membrane of live T cells. (A)** Schematic illustration of the sequential reaction strategy for CNP binding on the plasma membrane of live T cells. Chol-DNAs, assist-DNAs, and DNA NPs were sequentially applied to the cell. **(B)** Representative confocal images of Jurkat T cells bound with 72-CNPs, prepared by the sequential reaction. Bright-field (BF) (gray, left), 72-CNP (magenta, middle), and the merged images (right) are displayed. **(C)** Intensity distributions for 72-CNP-bound to Jurkat T cells, prepared by single and sequential reactions analyzed by flow cytometry. **(D-G)** Two sets of representative maximum intensity projection images for single-polarized (D), multi-polarized (E), unpolarized (F), and weak or unbound (G) Jurkat T cells incubated with 72-CNP for 30 min via the single reaction strategy. BF (gray, left), 72-CNP (magenta, middle), and the merged images (right) are shown. Scale bars in (B): 20 μm; in (D-G): 5 μm.

**Figure S3**

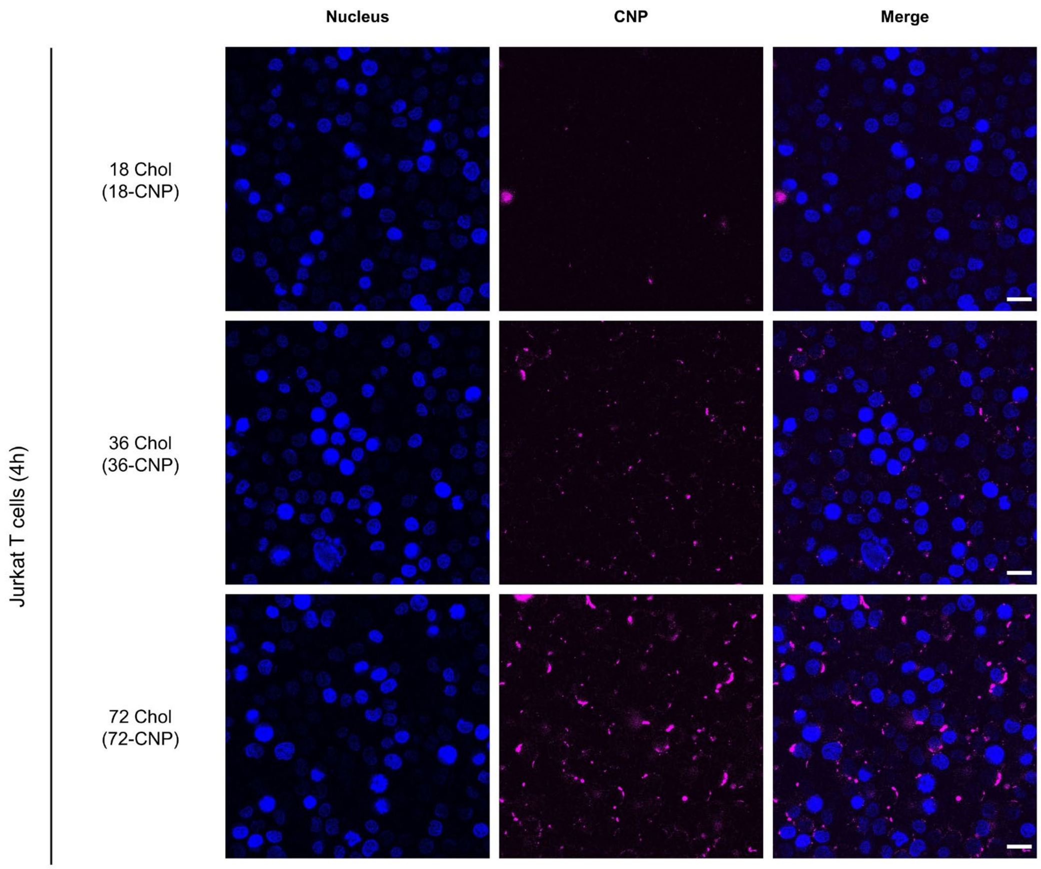


**Figure S3. CNP binding on the plasma membrane of live Jurkat T cell for 4 hours.** Representative confocal images of Jurkat T cells incubated 18-, 36-, 72-CNPs for 4 h. Images display nuclei stained with Hoechst (blue, left), CNP (magenta, middle), and the merged image (right) for each sample. Scale bars: 20 μm.

**Figure S4**

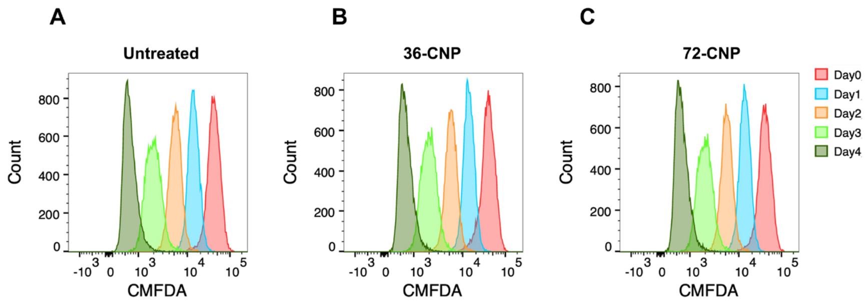


**Figure S4. Cell proliferation upon CNP binding for 4 days. (A-C)** Flow cytometry analysis was performed to assess Jurkat T cell proliferation over a 4-day period after CNP binding (day 0 to day 4). The cells were labeled with CMFDA dye and then incubated without treatment (A), or with 36-CNPs (B) or 72-CNPs (C) for 30 minutes on day 0.

**Figure S5**

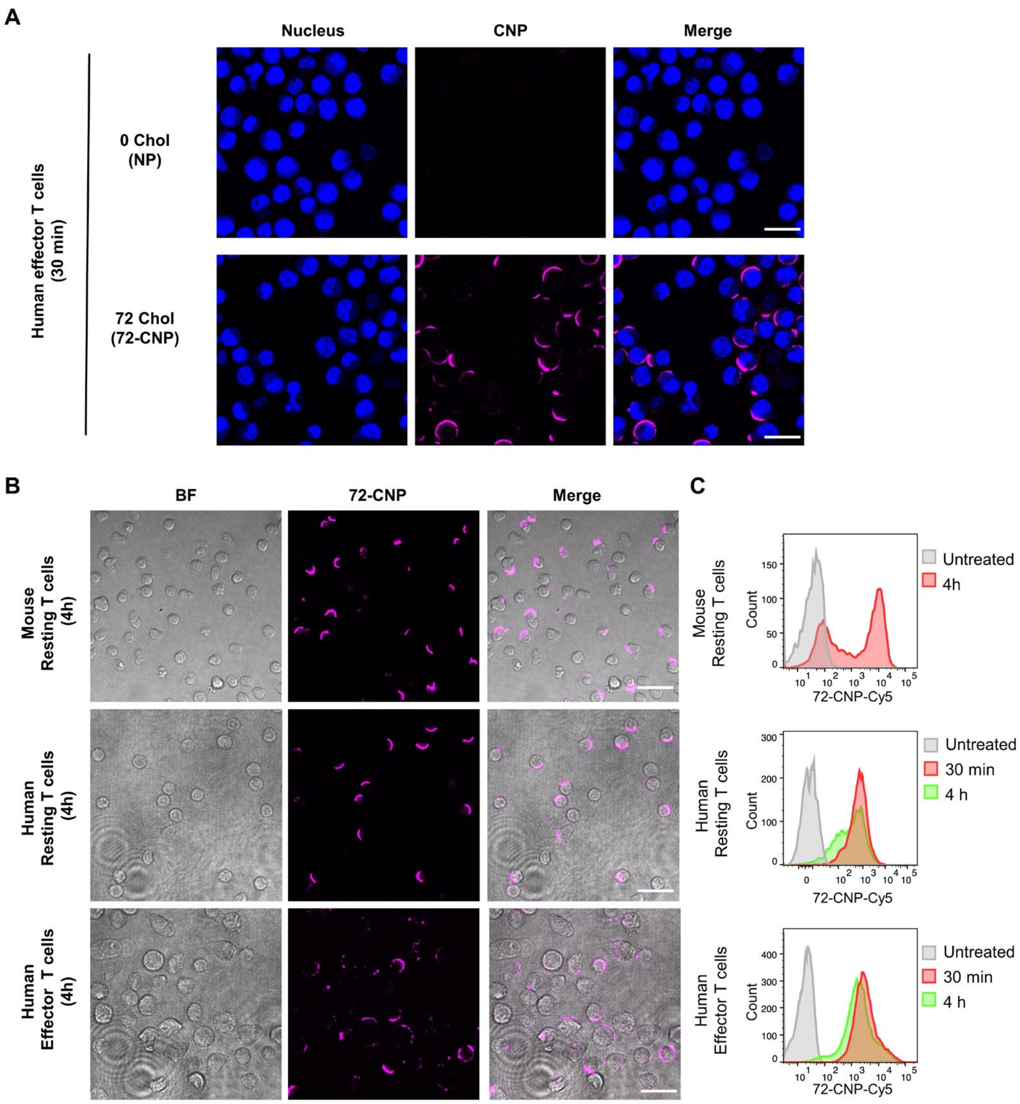


**Figure S5. Polarization of 72-CNPs on the plasma membrane of primary mouse and human T cells. (A)** Representative confocal images of effector human T cells incubated with DNA NP without cholesterol (0 Chol, upper), or 72-CNP (lower) for 30 min (scale bars: 20 μm). **(B)** Representative confocal images of mouse resting (top), human resting (middle), or human effector (bottom) T cells incubated with 72-CNP for 4 h. **(C)** The intensity distributions of the 72-CNP on the mouse resting (top), human resting (middle) or human effector (bottom) T cells incubated for 30 min (green) or 4 h (red) analyzed by flow cytometry. Scale bars in (B): 20 μm.

**Figure S6**

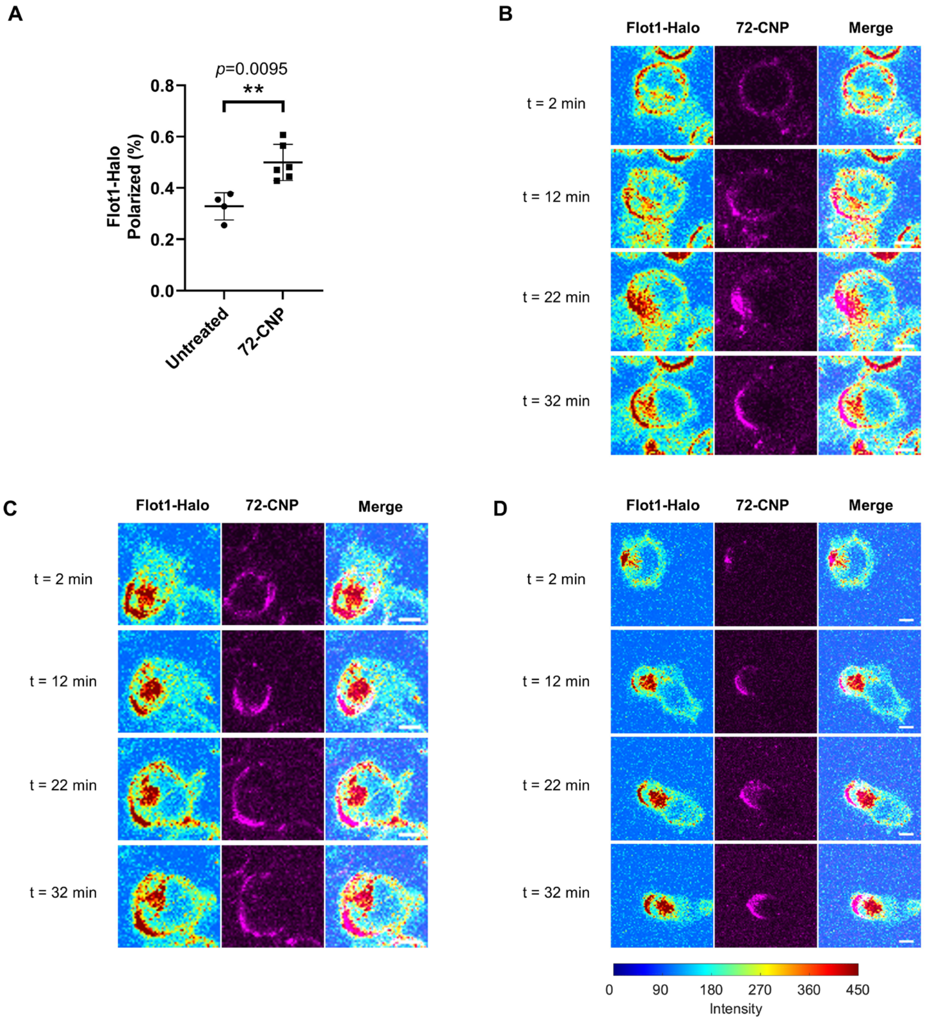


**Figure S6. CNP induces or enhances polarization of Flot1 on the T cell membrane. (A)** The percentage of polarized Flot-1 positive cells before (untreated, n= 234) and after incubating with 72-CNPs (n=143 cells) for 30 min, counted from the Flot1-Halo expressing Jurkat T cells within 212x212 $\mu m^2$ areas. Each dot represents the result counted from a single field of view collected from three independent experiments. The *p*-value (0.0095, **, $p \leq 0.01$) was calculated by a two-tailed unpaired Mann-Whitney test. Data in (A) are means ± SD. **(B-D)** Three representative cases of live time-lapse confocal images of Flot1-Halo-expressing Jurkat T cells from three independent experiments upon the addition of 72-CNPs are as follows: 72-CNPs initially bound homogenously to the polarized Flot1 negative cells and induced polarization of Flot1 (26/64 cells) (B); 72-CNPs initially bound homogenously to the polarized Flot1 positive cells, subsequently moving towards and enhancing the Flot1-polarization (21/64 cells) (C); 72-CNPs bound preferentially to the polarized Flot1 and enhanced its polarization (17/64 cells) (D). (See **Movie S1-S3**). The images were taken at 2, 12, 22 or 32 min after adding 72-CNPs. Flot1-Halo (color-coded (indicated at the bottom left), 72-CNP (magenta, middle) and the merge (right) of these images are presented. Scale bars: 5 μm.

**Figure S7**

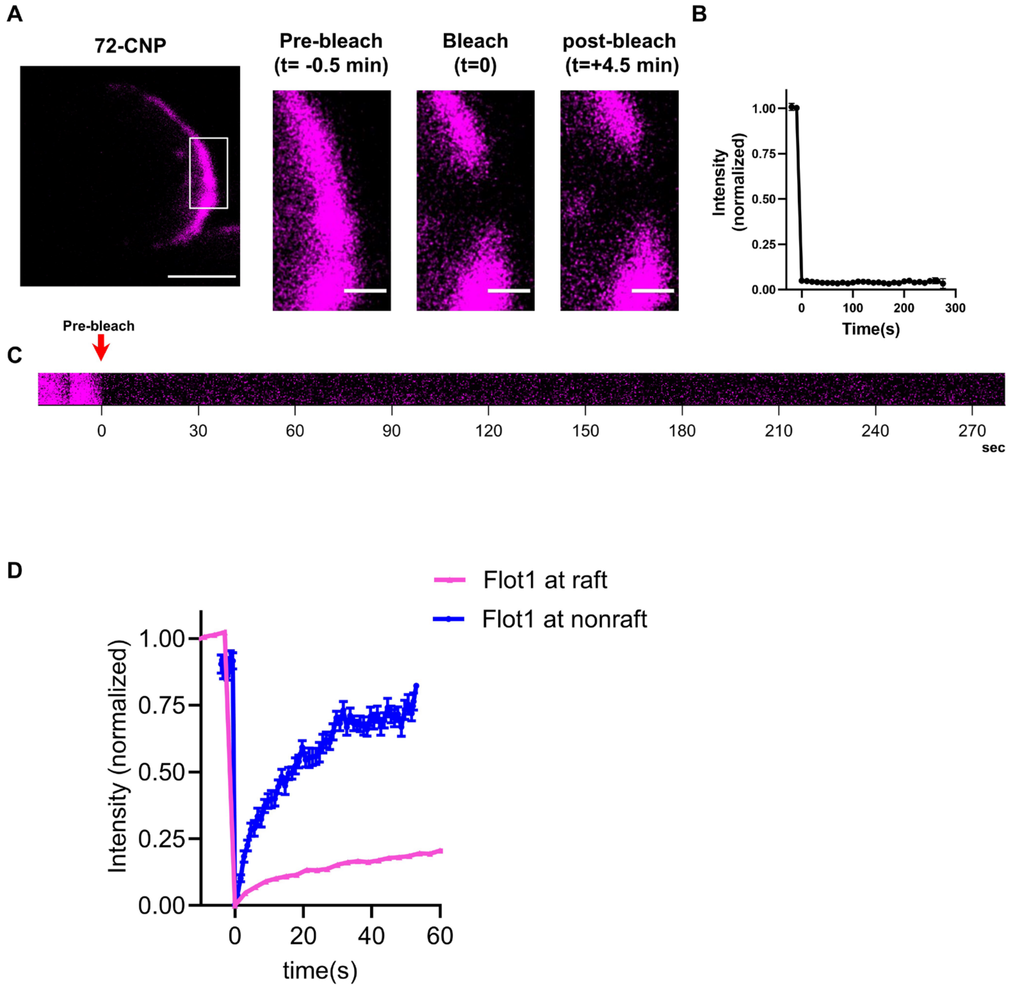


**Figure S7. FRAP analysis of the lateral mobility of 72-CNP bound on the Jurkat T cell membrane. (A)** Representative time-lapse confocal images of 72-CNP (magenta) in the area marked by a square in the left image taken before and after bleaching. Scale bars in the left image: 5µm; in the three right images: 1µm. The time points for each frame are indicated. **(B)** The mean FRAP tracings of 72-CNP measured for 5 minutes. (n = 15 cells from three independent experiments). **(C)** The filmstrip showing a fluorescence recovery of the bleached area of the cell in a marked in **Movie S4**. The bleached time is indicated with a red arrow. **(D)** The mean FRAP tracings of Flot1 on the 72-CNP bound cells measured for 1 minutes. Flot1 localized to the 72-CNP-bound (raft, blue line, n=26 cells) or unbound (non-raft, magenta line, n=14 cells) regions is measured. Data in (B) and (D) are mean ± SE. All data were collected from at least three independent experiments.

**Figure S8**

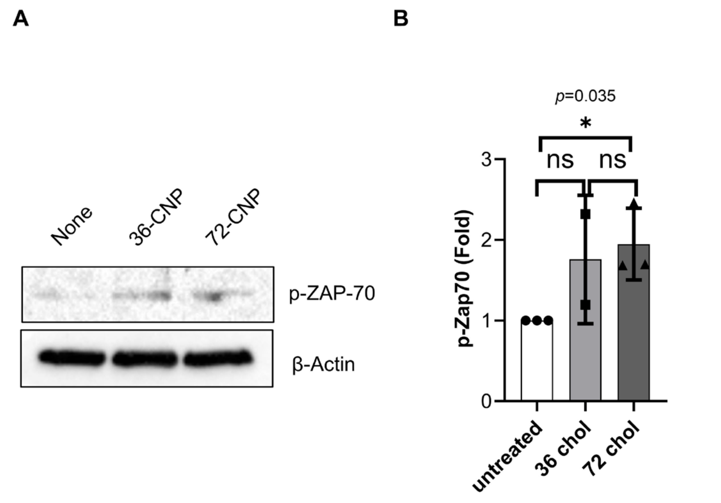


**Figure S8. CNPs induce or enhance early T cell activation signaling. (A-B)** Immunoblot analysis of phosphorylated ZAP-70 (p-ZAP-70) in human primary effector T cells incubated without or with 36- or 72-CNPs for 30 min. β-Actin levels represent the loading control. A representative image of immunoblotting (A) and quantification of p-ZAP-70 levels (B). The intensity of each p-ZAP-70 band relative to respective β-actin band was normalized to that of untreated samples. Each dot represents the band intensity quantified from an individual immunoblotting from three independent experiments. Data represent means ± SD.

**Movie S1.** A series of representative time-lapse confocal images of Flot1-Halo expressing Jurkat T cells show that 72-CNPs initially bound uniformly to the polarized Flot1-negative cell, subsequently inducing the formation of Flot1 polarization. Flot1-Halo (left), 72-CNP (magenta, middle), and the merged image (right) of these two are presented in each frame. The first frame was taken 2 min after adding the 72-CNPs to the sample and the movie was acquired for 30 min. Scale bars: 2 μm.

**Movie S2.** A series of representative time-lapse confocal images of Flot1-Halo expressing Jurkat T cells show that 72-CNPs initially bound uniformly, and then moved toward the existing polarized-Flot1. Images are similarly presented in **Movie S1**.

**Movie S3.** A series of representative time-lapse confocal images of Flot1-Halo expressing Jurkat T cells show that 72-CNPs preferentially bound to the polarized Flot1 and enhanced its polarization. Images are similarly presented in **Movie S1**.

**Movie S4.** A series of representative time-lapse confocal images during FRAP acquisition for 72-CNPs bound to the live T cell membrane. The bleached area is marked by a square. Scale bar: 1 μm.

**Movie S5.** A representative time-lapse series of confocal images during FRAP acquisition for Flot1-Halo (green, left) and 72-CNPs (magenta, right) on the live T cell membrane. The bleached areas are marked by a square for each channel. Scale bar: 1 μm.

**Table S1**

**Sequence of staple strands used in the DNA structures**

| Name | Sequence (5’ to 3’) |
|---|---|
| Chol-ssDNA | [Cholesterol] [TEG]GCCCAGACACGTTAACCAGAGGCTGACGTAGC |
| assist-DNA | CTCTGGTTAACGTGTCTGGGCTTTTTT |
| Fluor_DNA_01 | [Cyanine5] TTTTTGATGGTGGTTATTACCTTATGCGATTTTATTT |
| Fluor_DNA_02 | [Cyanine5] TTTAAGAAACGCAAAGACAAGGTGGCAACATATAATTT |
| Fluor_DNA_03 | [Cyanine5] TTTTTGCCGTTCCGGCAAAAGTTAAAGACGCAGAAACAGC |
| Fluor_DNA_04 | [Cyanine5] TTTACTACGTGAACCGCGTACTATGGTTGCTTTGTTT |
| Core_DNA_01 | ATTCTACAGAAGCCCCTTAGAGTGAATTTATCAAAGTTTGAA |
| Core_DNA_02 | ATCCCAACCTCCCGGTAGGAAAGTCCTGCAAAAGGTTAGTATTTTAATG |
| Core_DNA_03 | AGCCCGATAATTGCTTTTGCGAGATGATGGCAATTTCGGGAG |
| Core_DNA_04 | AAAAAAGCTAAAGTCAGTACCTGAGCAAAAGAAGATGTAAATCGTCGCT |
| Core_DNA_05 | TGTTTTAATTTGGGAAACAATGAATTATTCATTTCAATTTTC |
| Core_DNA_06 | ACCGGATTCTATCAGGGCGATGGCCCTTT |
| Core_DNA_07 | TTTAGTAATCTTGACAAGAGCTGACCAACGTGCGGTAATATCCAGAACAATATTT |
| Core_DNA_08 | TTTCGAGTAGATTTAGAGCCTCCTTTT |
| Core_DNA_09 | TTTCGGAATAGGTGTTACCGTAACACTTTT |
| Core_DNA_10 | TTTAAAGCCTCAGAGCGGCCAGAATTT |
| Core_DNA_11 | TTTCCGGAGACAGTCCAGCCAGCGTTT |
| Core_DNA_12 | TTTAATGGAAAGCGCGTTAATGCCCCCTTT |
| Core_DNA_13 | TTTATATGTACCCCGTTTCTTTGCTTT |
| Core_DNA_14 | TTTGTAGCCAGCTTTCTTTCGCACTTT |
| Core_DNA_15 | TTTATCGCCTGATAAATTGAACGGAGATTGCAAATCGCCATTAAAAATACCGTTT |
| Core_DNA_16 | TTTTTAGCGTCAGACGCCACCCTCAGATTT |
| Core_DNA_17 | TTTCGCACTCCAGCCCAGCACCGTTTT |
| Core_DNA_18 | TTTGGTTTTCCCAGTCATAACGGATTT |
| Core_DNA_19 | TTTCACGGAAAAAGACGATGCTGATTT |
| Core_DNA_20 | TTTCCCACAAGAATTGAGTAATATCAGAGAGATAATTT |
| Core_DNA_21 | TTTATCTTACCAACGCTAATACAATTTTATCCTGATTT |
| Core_DNA_22 | AAGAATGCGCAACCGGTCATTGCAGGCGCATCAACGTATCGGCCTCAGGAAGATTTT |
| Core_DNA_23 | GGTAATTGAGCGCTTAAGCCCGACAGAATCAAGTTTGCCTTTT |
| Core_DNA_24 | TTTACGTGCCGGACTTGTAACATCCTCACGACGGTGGTGAAGGGATAGCTCTTTT |
| Core_DNA_25 | TTTAGAACTGGCTCAGCTGATTGCTTT |
| Core_DNA_26 | TTTATCATTCCAAGAACGGTCGGCTGTCTTTCCTTTTT |
| Core_DNA_27 | TTTCGGTGGTGCCATCCCACCAACGGAGCTTTCTTAAGTTGGGTAACGCCAGTTT |
| Core_DNA_28 | TTTATTTAACAACGCCAACAATTGAGAATCGCCATTTT |
| Core_DNA_29 | AAACCAATCAATAAGTATTAAAAATCCTCATTAAAGCCAGTTT |
| Core_DNA_30 | TTTTCAATCCGCCGGGCGCGGTTGCGGTAATGGTTACACT |
| Core_DNA_31 | TTTCCAATCGCAAGACAAAAAATGCTGATGCAAATTTT |
| Core_DNA_32 | TTTTCGTCATAAACATCCCGTAAAGGGTTGATAATTCGCGTCTGGCCTTCCTTTT |
| Core_DNA_33 | TTTACCTTTTTTAATGGAACAATTTCATTTGAATTTTT |
| Core_DNA_34 | ATATAACTATATGTGAACGCGGGTTGATATAAGTATAGCCTTT |
| Core_DNA_35 | TTTGTGCCGGTGCCCCCTGTTACCTGAAATCACTAAAACTAGCATGTCAATCTTT |
| Core_DNA_36 | TTTACGTAAAACAGAAATATATCAAAATTATTTGCTTT |
| Core_DNA_37 | TTTTGCGGCGGGCCGTTTTGGTGCTGCATAAAGTTCAAAAGGGTGAGAAAGGTTT |
| Core_DNA_38 | TTTACTTTTTCATGAGGAAGCTTTGAGCAGAAGAATAGATAATACATTTGAGTTT |
| Core_DNA_39 | TTTGATTTAGAAGTACGATAGTTGTTT |
| Core_DNA_40 | TTTCACAGTTGAGGATCCCCGTCCGTGTTTGACATTAGCAAAATTAAGCAATTTT |

| | |
|---|---|
| Core_DNA_41 | TTTAACGAACCACCAGGACTAAAGTTT |
| Core_DNA_42 | TTTGCTCACAATTCCACACAACATACGCCTAAT |
| Core_DNA_43 | TTTTTACCGCCAGCCATTTGTATCTTT |
| Core_DNA_44 | TTTATTAATTGCGTTGCGCGAGTGAGGAGAATGGGAAGCAAACTCCAACAGGTTT |
| Core_DNA_45 | TCACTGCCTGCATTTTGCGTATGAGACGCTGAGAGCAGCAGGCGAAAATCCTGTTTT |
| Core_DNA_46 | TTTACGAGCACGTATTTCATCAAGTTT |
| Core_DNA_47 | TTTCGCGGGGAGAGGCGGTAATGAATTAACCCTCCCCTCAAATGCTTTAAACTTT |
| Core_DNA_48 | TTTCCTTCACCGCCTGGCCGGCAACATTATACCATTACGAGGCATAGTAAGATTT |
| Core_DNA_49 | TTTGCAACACTATCACGGCCAACGTTT |
| Core_DNA_50 | TTTCGCCGACAATGACAACTTGATACTTAGACTTTACAAAAGCCGTCATAAAAC |
| Core_DNA_51 | TTTAGTTCAGAAAACCTAACTCACTTT |
| Core_DNA_52 | TTTAGCGGAGTGAGAATAGAACTTTCAACAGTTTCTTT |
| Core_DNA_53 | TTTTCAGGATTAGAGTTGTTATCCTTT |
| Core_DNA_54 | AGTACCTTGATTCCCAATTCTGCGAATTT |
| Core_DNA_55 | CTTGCCCAACCGAAGACAGGAACTTCTTTGATTAGAAATGGATTATTTA |
| Core_DNA_56 | TTCAGTGTATTAAAGAACGTGGTAAAGCGTAACCA |
| Core_DNA_57 | ACCCAAAACAGACCTCAGAGCACTCAAACTATCGGAACGCTCATGGAAA |
| Core_DNA_58 | TTGAGATGTTAATATCAGTGAGGCCATTGTGAAAGGAAGGGA |
| Core_DNA_59 | GTAGTAAAGATAGGGTTGAGTGAGCTTGGCGGGCG |
| Core_DNA_60 | AGAACGAAACAACATCCTGAGCAAATTAACCGTTGAGATTCA |
| Core_DNA_61 | GACTCCAACGTCAAAGGGCGAGTTTTTTCTTAATG |
| Core_DNA_62 | ATCCCTTATAAATCAAAAGAACGAACGTGGCGACCACGCTGG |
| Core_DNA_63 | GTTGTTCCAGTTTGGAACAAGCTAAAGGAGTGTAG |
| Core_DNA_64 | CCACAGAATTTTCAAATAACCACCTCCGGCTTAGGAATATATTTTAGTT |
| Core_DNA_65 | AGGCAAGGGTTGTACCTGTGCCGATCCAGCGCAGTGTGATT |
| Core_DNA_66 | CATTAGATAACAGTTTAATTGCCAGACCACCATAATTGAATC |
| Core_DNA_67 | TCAGAACGGATTAGACCTAAACATATGCGTTATACATATAAAGTACCGA |
| Core_DNA_68 | TACTCAGTCGAGAGAGAAAACCAACAGTAGGGCTTATGTAATTTAGGCA |
| Core_DNA_69 | TTAAATGTGATAAAAGGCGTTAAATACGTTAACGGCATCAGAGGTGTGTTCAGCAA |
| Core_DNA_70 | TTTTTAGGGTAGCTATACCGATAGAAAAAGCCTGTTAAAGTA |
| Core_DNA_71 | TAAGAGGGGTCAGTCCAGTAATCCTAATTTACGAGTCATCGAGAACAAG |
| Core_DNA_72 | ATGAACGAAGCCCCGCGGCTGGTATGAGCCGGGTCACTCAGCTAA |
| Core_DNA_73 | ATCAGGTTGTAAACATTCTGTATCAACAATAGATATCATTAC |
| Core_DNA_74 | TACAGGAAGGAGGTTTTCATCACTTGCGGGAGGTT |
| Core_DNA_75 | GAATTTAAACAAATACCAAGTATTTTGCACCCAGCCGAGCGTCTTTCCA |
| Core_DNA_76 | TCATTTTGGATTCTCAGATATAGAAGTGTCCAGCATCAGCGGAGCTTACGGCTGGA |
| Core_DNA_77 | TTAAATTGGCGGATCGCGCCCGGCGTTTTAGCGAATCCAAAT |
| Core_DNA_78 | ATCAGAAGTAATCGCATCAATGTAAAGACTAAATCGCAAAGA |
| Core_DNA_79 | GAACCACCCAGAGCAGTTACAACACCCTGAACAAAGAAACAATGAAATACAGTATG |
| Core_DNA_80 | AGGGGACGCTTCTGCAACCGCAGCGTGGTGCTGGTCTAAATAGCA |
| Core_DNA_81 | TGGTGTAAGGCTGCAAGAAACACATAAAAACAGGGTAAGAAA |
| Core_DNA_82 | CGGAACGTCTTTGACGCTCAAATGGCTATTAGTCTAACACCGCCTGCAA |
| Core_DNA_83 | TAACGCCCAGACGAGTGCCAGCCGCTTTCCAGT |
| Core_DNA_84 | AGTTGAGAGAAGTTCCAGTCAACCTGAAAGCGTAATGAAAAA |
| Core_DNA_85 | TTGCCATCGGAAACACCGAAGTGATTAA |
| Core_DNA_86 | CGTTTTCCAGTAGCAATAATAAAATACA |
| Core_DNA_87 | GCCAGCTTCAGAGGAGTTACCAGAAGCGCTGGCAGCCTCCGGCGTTTTT |
| Core_DNA_88 | TGCGGGCGAACGGAAGTAAGCGAATACC |
| Core_DNA_89 | ATCACCAGGAAGGTCAATAGAAAATTCATATGGTT |
| Core_DNA_90 | TTGGGAAATTATTCATAAGTTTATTTTGACGTAGA |
| Core_DNA_91 | TTTCTCCTTGTAAAAAGGCGACGGCACCGACGACAATTAAATAAAAATA |
| Core_DNA_92 | GATAGACGGATCAAGTGATGAAGGGTAACGCGGTCCCAGAGCGAACGTC |
| Core_DNA_93 | CCAGTCCTCATTTGAAGCCGCACAGGCGGTGAAACCGAGGAA |
| Core_DNA_94 | AAACGTAGCGGTTGCATAAAAAAATCCCGTAAAAA |
| Core_DNA_95 | TGTACAAAAGGGCGACATTCATACCAGCGCCAAAGACATCGA |

| | |
|---|---|
| Core_DNA_96 | AGAGCAAGTCAGAGGAGCCTAAGTTGCTACCGCACCATGTAGGAGGCATCAACGCT |
| Core_DNA_97 | CAGCCATATTATTTTTAGACGGGAGAATCTATCTTGTCACCAATGAAAC |
| Core_DNA_98 | AAATAAATTGAAGCCAAGCCGTATCCCATAAGAGAAAATTCTAATTTCACTTTTTA |
| Core_DNA_99 | TCAGCAGGTGCCGGAAACCAG |
| Core_DNA_100 | GCCTTTACAGACGAACAATGGAGCCGCCACGG |
| Core_DNA_101 | CTTAAATCAAGATTATTTGCCCACCACCGGAACCG |
| Core_DNA_102 | AAGTGTTTAACGTCAAAAATGAGGGGGCTTATCCGGTATTCT |
| Core_DNA_103 | AACGCGAAATAGCACCTGTTTCCAGACGTAATTACCCGTGTG |
| Core_DNA_104 | TTGCCCTAAAAACAGGAAGAT |
| Core_DNA_105 | ACGACAATAAACAACATGTTGATAGAATAAACACCGGAATCA |
| Core_DNA_106 | AACAAGAAAAATAATTTTTATTGAGGCAGGTCAGA |
| Core_DNA_107 | TGCAGAACGCGAGCAAATCCGTGGGAACAAAC |
| Core_DNA_108 | TTTTTCATTGGGTTAATATATAACAAAAATTTTCATGGAAGGGTATTAACACTAAC |
| Core_DNA_109 | TGCCGGGCATCAGAACTCTGTCACGGTCTTCTTCGCGGGTACTTTCCTGTGTGAAA |
| Core_DNA_110 | TACCAGTATAAAGCTTTCGAGGCCTTGAGTAACAG |
| Core_DNA_111 | TTGCTTCTGATGAATGAATATTTGGATTCGTTATT |
| Core_DNA_112 | AAAACATAGCGATAGCTTACATGACCAAGTTACAAAATCGCG |
| Core_DNA_113 | ATCATAGATTAATTAATTACCTTTTACACATCAATGAGTAAC |
| Core_DNA_114 | GTCTGAGAGACTACTCTTCTGCGGGGTTTTGCTCA |
| Core_DNA_115 | AAGACGCTGAGAAGAATAAATATTAATGCCGGAGAG |
| Core_DNA_116 | GTCAATAATCCTTGCAGAGGCAACGGATGATTATCGAACAAAGGTCAGTCATCACC |
| Core_DNA_117 | TCGCCTGTTAGCTATATTTTC |
| Core_DNA_118 | ACAAACATCAAGAAGTGAGTGGGGATAGCAAGCCC |
| Core_DNA_119 | ATACCGGGGGTTTCCTGCGCGCCAAAAACATTATG |
| Core_DNA_120 | GAAAGAGGCGGGATGAGCCAGGAATTGAGGAAGGTAATTTTAAAAGTTT |
| Core_DNA_121 | GGTAAAAAGGGTAGAGGCGGTAACTAATAGATTAGCAATTCGACAACTC |
| Core_DNA_122 | AATACTGCTGACTAACCTCAAATATCGTGTAAAGCCTGGGGT |
| Core_DNA_123 | TGTTTAGAAAGCGGTCTAAAGTGGCAAATCAACAGATTATCA |
| Core_DNA_124 | GAAACCACCAGAAGGAGCGATAAAAACCCTCAATCAATATCT |
| Core_DNA_125 | ATAATCCTGATTGTACAGTAATTTGTCGTCTTTCC |
| Core_DNA_126 | GTTAGAACCTACCAAAGAAATGGATTTTGCTAAAC |
| Core_DNA_127 | CAGTATTTTAATGCTACCTACTAGAAGAGGGAGCTCGCCGCGGGGGTCGAGGTGCC |
| Core_DNA_128 | TTGAAAGCAGCAAAGAATACGCATTGGCTAGCAATACGGTACGCCAGAA |
| Core_DNA_129 | TATCTAACAGTGCCTTTTTGATCGTCTGTAATAACTTAAAGGGCTGCGC |
| Core_DNA_130 | AATATCTTTAGGAGATCCTTTATCAGCTTGCTTTC |
| Core_DNA_131 | GAGCCGGAAGCATACATGGTCTGATAAGAGGTCAT |
| Core_DNA_132 | ACCTGTCCGATAAAAACCAAA |
| Core_DNA_133 | AGTAATAAAAGGGACATTCAAGGCCGAGTAAAAGAGTCTGTC |
| Core_DNA_134 | TGGCACAGACAATAACGCTGACGTCACCCTCAGCA |
| Core_DNA_135 | GCGAACTGATAGCCAGAGGTGCAACGGCTACAGAG |
| Core_DNA_136 | CTAAAACCAGGAAACCTTGCTTTTCCTCACAGGGC |
| Core_DNA_137 | CGGGTGGCCAACAGAGATAGAACTTGCTGATTATAGTCAGAAGC |
| Core_DNA_138 | CCTTCTGCACGACCCATCACGAAGTGTTGAAAGGA |
| Core_DNA_139 | GCCTGAGATTTTGACCCCCAGCGATTAT |
| Core_DNA_140 | CGTTTACAAAAGGAAGTCAGGATTGTGACCGAAATCGGCAAATTTGCCCAGTTGCA |
| Core_DNA_141 | CAAAAGAACTGGCACCCTTTTAAGCGCA |
| Core_DNA_142 | GACTCCTTATTACGGCAATAGTAACTGA |
| Core_DNA_143 | TTAGCAATCACAATAAATATTGACGGAA |
| Core_DNA_144 | TACATAACCACGGAATTAAAGGTGAATTATCACTTT |
| Core_DNA_145 | TCGTCTCGCCTTTAACTTAAATTTCTGC |
| Core_DNA_146 | ACGCAATAATAACGAGATAGCGAGAATAGATTTTT |
| Core_DNA_147 | CTGAGGCTCGGTTTGCCCGAAATACTTCTGAATAAGGTTTAACGTCAGA |
| Core_DNA_148 | AAGCGAACTCCTTTATAGCTGCGAGCTCGAATTCGTAGAATTATCATCATATTCCT |
| Core_DNA_149 | CTAGGGCGCTGGCAGAGCCCCCGATTTA |
| Core_DNA_150 | CGGTCACGATTTTACTGACCAACTTTGA |

| | |
|---|---|
| Core_DNA_151 | CCACACCAAACAGGAGGCCGAATCACTT |
| Core_DNA_152 | CGCCGCTGTTAGAAAGGCGCATAGGCTG |
| Core_DNA_153 | GCAAGCGTTTCTTTTCACCAGTTGGGCGCCAGGGT |
| Core_DNA_154 | AGAAAGCTTTATAAAAACGAACTAACGG |
| Core_DNA_155 | AGGAATTCTGTATGTGCGTAGTTAATTACATTTAAACAGTACATAAATC |
| Core_DNA_156 | GTACGGTAACCTGTATTGCTTTGAATCCAGCACGCGTGCCTG |
| No_Chol_anchor_01x1 | CTAATGCAATCTACGGTTTAATTTCAAC |
| No_Chol_anchor_01x2 | ACGGGGAAAGCCGGTAGCCCGATTGGGC |
| No_Chol_anchor_01x3 | TTATTACAGGTAGGTGACGAGAAACACC |
| No_Chol_anchor_01x4 | ACTAAATCGGAACCAGTCCACAATAAGG |
| No_Chol_anchor_01x5 | CCATGTTACGGTGTTCAACGTAACAAAG |
| No_Chol_anchor_01x6 | ATCACCCAAATCAAAAAACCGATTCATT |
| No_Chol_anchor_02x1 | TTTAATCACGTTGGGAAGAAAAGATACA |
| No_Chol_anchor_02x2 | ACTGGATTTGCAAAATTTAGGAATACCA |
| No_Chol_anchor_02x3 | AATCATAAGGAAAGATTCATC |
| No_Chol_anchor_02x4 | CACCAACACACTCAAGGCGCAGACGGTC |
| No_Chol_anchor_02x5 | CTGCTCAAAGAGGACAGATGAACTTAGC |
| No_Chol_anchor_02x6 | GTTTCCAAAAGTACTGTCGAAATCCGCG |
| No_Chol_anchor_03x1 | TAATTCGCTTTACCCGGAATCGTCATAA |
| No_Chol_anchor_03x2 | CATTCAAATAGCGAGAGGCTTAGCGTCC |
| No_Chol_anchor_03x3 | ATTGCATCAAAATTGCGTAATAGTAAAA |
| No_Chol_anchor_03x4 | TTGCCAGAGGGGAAAAGAATACACTAAACTAAAAC |
| No_Chol_anchor_03x5 | TATATCGGAACGTACGTAATGCCACT |
| No_Chol_anchor_03x6 | ACCTGCTACCAAGCGCGAAACTTAAACG |
| No_Chol_anchor_04x1 | ATATTCAATCAAAAATCAGGTAGCTTCA |
| No_Chol_anchor_04x2 | AATATGCTAGAGCTAAGACTTCAAATAT |
| No_Chol_anchor_04x3 | AGGCCGCTTAGATTAAGAGGA |
| No_Chol_anchor_04x4 | TTGAAAAAATTGTATTGCAGGGAGTTAA |
| No_Chol_anchor_04x5 | ACGAAGGGCGAAAGACAGCATTCGGTCG |
| No_Chol_anchor_04x6 | AAAGGAAAAACAGCAACCATCGCCCACG |
| No_Chol_anchor_05x1 | CCAATAAGGTCAATGTCTGGAAGTTTCA |
| No_Chol_anchor_05x2 | CGCGTTTTTTTGCGGATGGCTAACTAAA |
| No_Chol_anchor_05x3 | GCGCGAGCTGAAATGCTGTAGCTCAACA |
| No_Chol_anchor_05x4 | TGAATATAATGCTCCAAAAGGAGCCTTTTCTCCAA |
| No_Chol_anchor_05x5 | ACGCCTGGAATTTTGCGAATAATAATTT |
| No_Chol_anchor_05x6 | CATAACCGAGGTGAATTTCTTCAACTAA |
| No_Chol_anchor_06x1 | TTCCATATACATTTCGCAAATATCATAC |
| No_Chol_anchor_06x2 | AACCCTCTTGCGGGTAATAGTAGTAGCA |
| No_Chol_anchor_06x3 | AGCGTAACGAAGGTGGCATCA |
| No_Chol_anchor_06x4 | CAGAACCCACCCTCCAGCCCTCATAGTT |
| No_Chol_anchor_06x5 | TTTCACGAGACGTTAGTAAATTAGCATT |
| No_Chol_anchor_06x6 | TTTGAGTTTCGTCACCAGTACA |
| No_Chol_anchor_07x1 | AAACAAGTTCTAGCCAATGCCTGAGTAA |
| No_Chol_anchor_07x2 | TTAACATACCCTGTAATACTTATATATT |
| No_Chol_anchor_07x3 | ATTTTTGAGAGATACGCAAGGATAAAAA |
| No_Chol_anchor_07x4 | TTTATTTCAACGCCACCCTCAGAGCCACGCCACCC |
| No_Chol_anchor_07x5 | AAACATGAGTGCCGGAGGTTTAGTACCG |
| No_Chol_anchor_07x6 | AACTACAAATAGGAACCCATGATCACCG |
| No_Chol_anchor_08x1 | TGTGTAGATGATATTCAACCGAGAATCG |
| No_Chol_anchor_08x2 | TTTGTTATTTAAATCATTGCCTGAGAGT |
| No_Chol_anchor_08x3 | GAGAAGGATTCTACAAAGGCT |
| No_Chol_anchor_08x4 | ATGGCTTTTAACGGCTGAGACTCCTCAA |
| No_Chol_anchor_08x5 | CCACCCTGTACCAGGCGGATAAAAGTAT |
| No_Chol_anchor_08x6 | TTTTGCCTATTTCGGAACCTAT |
| No_Chol_anchor_09x1 | CATCTGCACCCGTCTTAACCAATAGGAA |

| | |
|---|---|
| No_Chol_anchor_09x2 | CTGGAGCTGTATAAGCAAATAAATCAGC |
| No_Chol_anchor_09x3 | TGACCGTAATGGACGTTTAAAATTCGCA |
| No_Chol_anchor_09x4 | GTTAATATTTTGGTACTGGTAATAAGTTTTGATGA |
| No_Chol_anchor_09x5 | CAGAGCCATTCACACCGTTCCAGTAAGC |
| No_Chol_anchor_09x6 | TATTCTGTGCCCGTATAAACAAGTCTCT |
| No_Chol_anchor_10x1 | CGCCATCGTGAGCGAGTAACACAGTTTG |
| No_Chol_anchor_10x2 | CTCTTCGGCCATTCGATGGGCGCATCGT |
| No_Chol_anchor_10x3 | CCAGCATTGGATAGGTCACGT |
| No_Chol_anchor_10x4 | CCCCTTAACCGGAACACCAGAGCCGCCG |
| No_Chol_anchor_10x5 | GTCATACCGATTGGCCTTGATGCCACCA |
| No_Chol_anchor_10x6 | TTTACCGCCACCCTCAGAGCCA |
| No_Chol_anchor_11x1 | CGGAATTCAAGCTTGGCGAAAGGGGGAT |
| No_Chol_anchor_11x2 | AACCGTGGCAAAGCGCCATTCCTATTAC |
| No_Chol_anchor_11x3 | TAACCTCACCGGGCCTAAGGGCGATCGG |
| No_Chol_anchor_11x4 | GCAACTGTTGGGTTTCATAATCAAAATCTTAGCGT |
| No_Chol_anchor_11x5 | TTAGAGCCCGTAATATCGGCATTTTCGG |
| No_Chol_anchor_11x6 | CCACCCTCCTCCCTCAGAGCCTGTAGCG |
| No_Chol_anchor_12x1 | GTGCTGCACGACGGCCAGTGCTGTGAGA |
| No_Chol_anchor_12x2 | CCGCCAGCAGTTGGCAGCGCCATGTTTA |
| No_Chol_anchor_12x3 | ATTAGCAAGAAACAATCGGCG |
| No_Chol_anchor_12x4 | ACCGATTGAGGGAGGTAGCACCATTACC |
| No_Chol_anchor_12x5 | TCATAGCCATCGATAGCAGCACAGCAAA |
| No_Chol_anchor_12x6 | TTTCGTCACCGACTTGAGCCAT |
| Chol_anchor_01x1 | CTAATGCAATCTACGGTTTAATTTCAACGCTACGTCAGC |
| Chol_anchor_01x2 | ACGGGGAAAGCCGGTAGCCCGATTGGGCGCTACGTCAGC |
| Chol_anchor_01x3 | TTATTACAGGTAGGTGACGAGAAACACCGCTACGTCAGC |
| Chol_anchor_01x4 | ACTAAATCGGAACCAGTCCACAATAAGGGCTACGTCAGC |
| Chol_anchor_01x5 | CCATGTTACGGTGTTCAACGTAACAAAGGCTACGTCAGC |
| Chol_anchor_01x6 | ATCACCCAAATCAAAAAACCGATTCATTGCTACGTCAGC |
| Chol_anchor_02x1 | TTTAATCACGTTGGGAAGAAAAGATACAGCTACGTCAGC |
| Chol_anchor_02x2 | ACTGGATTTGCAAAATTTAGGAATACCAGCTACGTCAGC |
| Chol_anchor_02x3 | AATCATAAGGAAAGATTCATCGCTACGTCAGC |
| Chol_anchor_02x4 | CACCAACACACTCAAGGCGCAGACGGTCGCTACGTCAGC |
| Chol_anchor_02x5 | CTGCTCAAAGAGGACAGATGAACTTAGCGCTACGTCAGC |
| Chol_anchor_02x6 | GTTTCCAAAAGTACTGTCGAAATCCGCGGCTACGTCAGC |
| Chol_anchor_03x1 | TAATTCGCTTTACCCGGAATCGTCATAAGCTACGTCAGC |
| Chol_anchor_03x2 | CATTCAAATAGCGAGAGGCTTAGCGTCCGCTACGTCAGC |
| Chol_anchor_03x3 | ATTGCATCAAAATTGCGTAATAGTAAAAGCTACGTCAGC |
| Chol_anchor_03x4 | TTGCCAGAGGGGAAAAGAATACACTAAACTAAAACGCTACGTCAGC |
| Chol_anchor_03x5 | GATATATCGGAACGTACGTAATGCCACTGCTACGTCAGC |
| Chol_anchor_03x6 | ACCTGCTACCAAGCGCGAAACTTAAACGGCTACGTCAGC |
| Chol_anchor_04x1 | ATATTCAATCAAAAATCAGGTAGCTTCAGCTACGTCAGC |
| Chol_anchor_04x2 | AATATGCTAGAGCTAAGACTTCAAATATGCTACGTCAGC |
| Chol_anchor_04x3 | AGGCCGCTTAGATTAAGAGGAGCTACGTCAGC |
| Chol_anchor_04x4 | TTGAAAAAATTGTATTGCAGGGAGTTAAGCTACGTCAGC |
| Chol_anchor_04x5 | ACGAAGGGCGAAAGACAGCATTCGGTCGGCTACGTCAGC |
| Chol_anchor_04x6 | AAAGGAAAAACAGCAACCATCGCCCACGGCTACGTCAGC |
| Chol_anchor_05x1 | CCAATAAGGTCAATGTCTGGAAGTTTCAGCTACGTCAGC |
| Chol_anchor_05x2 | CGCGTTTTTTTGCGGATGGCTAACTAAAGCTACGTCAGC |
| Chol_anchor_05x3 | GCGCGAGCTGAAATGCTGTAGCTCAACAGCTACGTCAGC |
| Chol_anchor_05x4 | TGAATATAATGCTCCAAAAGGAGCCTTTTCTCCAAGCTACGTCAGC |
| Chol_anchor_05x5 | ACGCCTGGAATTTTGCGAATAATAATTTGCTACGTCAGC |
| Chol_anchor_05x6 | CATAACCGAGGTGAATTTCTTCAACTAAGCTACGTCAGC |
| Chol_anchor_06x1 | TTCCATATACATTTCGCAAATATCATACGCTACGTCAGC |
| Chol_anchor_06x2 | AACCCTCTTGCGGGTAATAGTAGTAGCAGCTACGTCAGC |

| | |
|---|---|
| Chol_anchor_06x3 | AGCGTAACGAAGGTGGCATCAGCTACGTCAGC |
| Chol_anchor_06x4 | CAGAACCCACCCTCCAGCCCTCATAGTTGCTACGTCAGC |
| Chol_anchor_06x5 | TTTCACGAGACGTTAGTAAATTAGCATTGCTACGTCAGC |
| Chol_anchor_06x6 | TTTGAGTTTCGTCACCAGTACAGCTACGTCAGC |
| Chol_anchor_07x1 | AAACAAGTTCTAGCCAATGCCTGAGTAAGCTACGTCAGC |
| Chol_anchor_07x2 | TTAACATACCCTGTAATACTTATATATTGCTACGTCAGC |
| Chol_anchor_07x3 | ATTTTTGAGAGATACGCAAGGATAAAAAGCTACGTCAGC |
| Chol_anchor_07x4 | TTTATTTCAACGCCACCCTCAGAGCCACGCCACCCGCTACGTCAGC |
| Chol_anchor_07x5 | AAACATGAGTGCCGGAGGTTTAGTACCGGCTACGTCAGC |
| Chol_anchor_07x6 | AACTACAAATAGGAACCCATGATCACCGGCTACGTCAGC |
| Chol_anchor_08x1 | TGTGTAGATGATATTCAACCGAGAATCGGCTACGTCAGC |
| Chol_anchor_08x2 | TTTGTTATTTAAATCATTGCCTGAGAGTGCTACGTCAGC |
| Chol_anchor_08x3 | GAGAAGGATTCTACAAAGGCTGCTACGTCAGC |
| Chol_anchor_08x4 | ATGGCTTTTAACGGCTGAGACTCCTCAAGCTACGTCAGC |
| Chol_anchor_08x5 | CCACCCTGTACCAGGCGGATAAAAGTATGCTACGTCAGC |
| Chol_anchor_08x6 | TTTTGCCTATTTCGGAACCTATGCTACGTCAGC |
| Chol_anchor_09x1 | CATCTGCACCCGTCTTAACCAATAGGAAGCTACGTCAGC |
| Chol_anchor_09x2 | CTGGAGCTGTATAAGCAAATAAATCAGCGCTACGTCAGC |
| Chol_anchor_09x3 | TGACCGTAATGGACGTTTAAAATTCGCAGCTACGTCAGC |
| Chol_anchor_09x4 | GTTAATATTTTGGTACTGGTAATAAGTTTTGATGAGCTACGTCAGC |
| Chol_anchor_09x5 | CAGAGCCATTCACACCGTTCCAGTAAGCGCTACGTCAGC |
| Chol_anchor_09x6 | TATTCTGTGCCCGTATAAACAAGTCTCTGCTACGTCAGC |
| Chol_anchor_10x1 | CGCCATCGTGAGCGAGTAACACAGTTTGGCTACGTCAGC |
| Chol_anchor_10x2 | CTCTTCGGCCATTCGATGGGCGCATCGTGCTACGTCAGC |
| Chol_anchor_10x3 | CCAGCATTGGATAGGTCACGTGCTACGTCAGC |
| Chol_anchor_10x4 | CCCCTTAACCGGAACACCAGAGCCGCCGGCTACGTCAGC |
| Chol_anchor_10x5 | GTCATACCGATTGGCCTTGATGCCACCAGCTACGTCAGC |
| Chol_anchor_10x6 | TTTACCGCCACCCTCAGAGCCAGCTACGTCAGC |
| Chol_anchor_11x1 | CGGAATTCAAGCTTGGCGAAAGGGGGATGCTACGTCAGC |
| Chol_anchor_11x2 | AACCGTGGCAAAGCGCCATTCCTATTACGCTACGTCAGC |
| Chol_anchor_11x3 | TAACCTCACCGGGCCTAAGGGCGATCGGGCTACGTCAGC |
| Chol_anchor_11x4 | GCAACTGTTGGGTTTCATAATCAAAATCTTAGCGTGCTACGTCAGC |
| Chol_anchor_11x5 | TTAGAGCCCGTAATATCGGCATTTTCGGGCTACGTCAGC |
| Chol_anchor_11x6 | CCACCCTCCTCCCTCAGAGCCTGTAGCGGCTACGTCAGC |
| Chol_anchor_12x1 | GTGCTGCACGACGGCCAGTGCTGTGAGAGCTACGTCAGC |
| Chol_anchor_12x2 | CCGCCAGCAGTTGGCAGCGCCATGTTTAGCTACGTCAGC |
| Chol_anchor_12x3 | ATTAGCAAGAAACAATCGGCGGCTACGTCAGC |
| Chol_anchor_12x4 | ACCGATTGAGGGAGGTAGCACCATTACCGCTACGTCAGC |
| Chol_anchor_12x5 | TCATAGCCATCGATAGCAGCACAGCAAAGCTACGTCAGC |
| Chol_anchor_12x6 | TTTCGTCACCGACTTGAGCCATGCTACGTCAGC |
